# Mild blast TBI raises gamma connectivity, EEG power, and reduces GABA interneuron density

**DOI:** 10.1101/2023.12.01.569541

**Authors:** Hazel G May, Konstantinos Tsikonofilos, Cornelius K Donat, Magdalena Sastre, Andriy S Kozlov, David J Sharp, Michael Bruyns-Haylett

**Affiliations:** Department of Brain Sciences, Imperial College London, W12 0NN London, United; Department of Bioengineering, Imperial College London, SW7 2AZ London, United Kingdom; Helmholtz-Zentrum Dresden-Rossendorf, Institute of Radiopharmaceutical Cancer Research, Bautzner Landstraße 400, 01328 Dresden, Germany

## Abstract

At least one traumatic brain injury (TBI) will be experienced by approximately 50-60 million of the world’s population in their lifetime and is the biggest cause of death and disability in those under 40. Mild traumatic brain injury (mTBI) can induce subtle changes but have long-lasting effects that may be difficult to detect through conventional neurological assessment, including standard clinical imaging techniques. These changes can lead to an increased risk of future neurodegeneration and emphasises the need to use more sensitive diagnostic tools such as EEG in order to identify injury and opportunities for therapeutic intervention.

In this study, we investigated electrophysiological and histopathological changes in a rat model of mild blast-induced TBI. We used a 32-channel EEG electrode array to detect global and local changes in neural activity and functional connectivity in acute (3 to 4-hours) as well as chronic phases (1 and 3-months) post-injury. GABAergic inhibitory interneurons, crucial for maintaining an excitatory/inhibitory balance, were quantified using immunohistochemistry.

Mild blast-induced TBI had minimal effects on resting power and connectivity at the acute timepoint but resulted in resting-state global power increases at all frequencies as well as a relative power increase in slow-wave frequencies in the chronic phase post-injury. Functional connectivity increases in the gamma frequency along with increases in power in the chronic phase pointed towards an alteration in the excitatory/inhibitory balance. Indeed, electrophysiological changes were associated with reduced density of GABAergic interneurons at 7-days, 1-month, and 3months post-injury, with a decrease in somatostatin-positive cell density in the 5th layer of all cortical regions of interest, and a parvalbumin decrease in the 5^th^ layer of the primary auditory cortex. In contrast, the total number of neurons, measured by NeuN did not change significantly, thus demonstrating a biased impact on inhibitory interneuron populations.

Our work demonstrates that the techniques and metrics of injury assessment employed in this study are sensitive enough to reflect the subtle changes present in mTBI and therefore hold potential clinical relevance. By using non-invasive EEG assessments and histopathology, we were able to reveal direct correlates and potential sources of the abnormalities caused by mild blast-induced TBI.

## Introduction

Recent publicity surrounding the development of neurodegenerative disease in military veterans and high-profile sports personalities has sparked a surge in Traumatic Brain Injury (TBI) research (Jordan 2000; Omalu et al. 2005; Goldstein et al. 2012; Mckee and Robinson 2014; Stewart et al. 2016; Mez et al. 2017; Mackay et al. 2019; Zimmerman et al. 2021). This has drawn attention to the potential long-term effects of TBI, specifically regarding the increased risk of developing conditions such as Chronic Traumatic Encephalopathy, Alzheimer’s disease and other forms of dementia (Plassman et al. 2000; Barnes et al. 2014), with multiple TBI lifetime events also a key risk factor for future neurodegeneration (Mckee and Robinson 2014).

In the context of modern warfare, blast-induced traumatic brain injury (bTBI) is a common outcome for service personnel (Taber, Warden, and Hurley 2006; Marshall et al. 2012). An estimated 10-20% of soldiers deployed in the Afghanistan and Iraq experienced TBI (Elder et al. 2010; Elder and Cristian 2009), and TBI was considered a signature injury of these conflicts (Elder and Cristian 2009) with 77% of TBI cases of United states soldiers classified as mild TBI (mTBI) (Marshall et al. 2012). Part of this rise in incidence can be explained by the increased use of improvised explosive devices (IEDs), especially in asymmetric conflicts, such as Iraq and Afghanistan (Hoge et al. 2008; Elder et al. 2010). In addition, the use of modern body armour and helmets along with advancements in battlefield medicine have resulted in increased survival rates for what were once previously fatal injuries, (Elder et al. 2010; Magnuson, Leonessa, and Ling 2012).

A big challenge in the management of TBI, especially mild TBI (concussion), is that symptoms often go unnoticed and unreported by sufferers (Hoge et al. 2008; Frost et al. 2013; Robinson et al. 2015; Tenovuo et al. 2021). And even if reported, TBI can still escape typical clinical assessment measures (Prince and Bruhns 2017; Maas et al. 2022). Indeed, bTBI is typically detected through a neurological assessment followed by brain imaging such as computerised tomography (CT) or Magnetic Resonance Imaging (MRI). However, this diagnostic protocol does not always detect milder forms of brain injury (Shin et al. 2017), as macroscopic changes spotted on structural scans in more severe forms of bTBI are rarely apparent in mild bTBI. Importantly, if no injury is identified then no recovery period will be mandated (Tenovuo et al. 2021; Maas et al. 2022). This is concerning because repeated TBI events can result in more severe and long-term injury if a recovery period is not observed (Meehan et al. 2012). These challenges highlight the urgency to find an effective method of identifying the initial occurrence of even a mild TBI to manage injury and recovery effectively. Therefore, diagnosis of mild TBI calls for more sensitive techniques such as measuring changes in neural activity using EEG or MEG to provide a functional snapshot of brain-wide activity potentially sensitive enough to identify changes in the absence of traditional TBI imaging markers of pathology.

By using high temporal resolution functional imaging techniques such as EEG and MEG, it is possible to identify changes in the absence of traditional TBI imaging markers of pathology. For example, an MEG approach (Huang et al. 2012; Huang et al. 2014) outperformed a structural MRI diagnosis tool, where they demonstrated that 87% of both blast and non-blast mTBI participants had the typical TBI hallmark feature of abnormally high delta (1-4 Hz) power relative to controls (Gloor, Ball, and Schaul 1977; Ball, Gloor, and Schaul 1977; Lewine et al. 1999; Huang et al.

2009; Franke et al. 2016; Lewine et al. 2019; Buchanan, Ros, and Nahas 2021; Mallas et al. 2022). This is in stark contrast to only 20% of the mTBI cohort showing any abnormalities in MRI. In addition, Franke et al. ^(Franke^ ^et^ ^al. 2016)^ used EEG in bTBI and non-bTBI cohorts, and also reported increased delta and theta power changes as a lasting chronic effect of bTBI in the absence of clinically assessed symptoms. Power spectrum changes are not the only electrophysiological abnormality seen in TBI. Functional connectivity (FC) changes, quantified by the statistical dependency of signals from different brain regions (Bowyer 2016; Sharp, Scott, and Leech 2014), are also observed (Lennon et al. 2023; Wang, Ethridge, et al. 2017), resulting in the altered ability of brain-wide networks to form and transmit information. The finding of electrophysiological correlates of pathology in populations who show no clinical signs of injury is important, as those who have experienced one injury have a higher predisposition for repeated injury and therefore a potential increased chance of developing future neurodegenerative disease (Lennon et al. 2023).

White matter damage has been proposed as the source of power and FC changes in EEG caused by TBI (Gloor, Ball, and Schaul 1977; Wang, Costanzo, et al. 2017). However, this is not the only cellular change that can affect network dynamics, and there has been little investigation of the more subtle structural and functional changes to the delicate balance between the excitatory and inhibitory neurons integral to the brain’s homeostasis. Known as the excitation/inhibition balance (E/I), it plays an important role in normal neural function and an imbalance can result in neurological disorders and seizures (Ferguson and Gao 2018). Inhibitory GABAergic interneurons play a crucial role in the E/I balance, and this extends to brain injury where two subtypes of inhibitory interneurons (expressing parvalbumin or somatostatin) have been found to be affected by TBI (Lowenstein et al. 1992; Toth et al. 1997; Hunt, Scheff, and Smith 2011; Cantu et al. 2015; Nichols et al. 2018; Frankowski, Kim, and Hunt 2019). Parvalbumin expressing (PV) interneurons are the most abundant cortical interneurons (30-40% of interneurons) (Scholl et al. 2015; Tremblay, Lee, and Rudy 2016) whereas somatostatin expressing (SST) interneurons comprise ∼30% of interneuron (IN) density (Huusko et al. 2015; Tremblay, Lee, and Rudy 2016). Given that it has been shown that perturbations to an interneuron population, firing pattern, or alteration in Ca^2+^-binding protein (e.g., PV) density can cause the E/I balance to be affected and result in behavioural as well as functional deficits (Sohal et al. 2009; Cantu et al. 2015; Filice et al. 2016; Ferguson and Gao 2018; Vascak et al. 2018; Sohal and Rubenstein 2019), it is important to explore how different types of TBI affect the E/I balance. In addition, despite the prevalence of mild bTBI, it is unknown whether the E/I pathologies and possible impact on network connectivity match that reported in blunt impact TBI (Cantu et al. 2015; Vascak et al. 2018).

To further examine the potential consequences of alterations in inhibitory function on brain activity, a preclinical model is necessary. In this research, we explored the potential value of EEG as a complementary addition to the existing suite of diagnostic tools within the context of mild bTBI. High-density EEG was used to measure the functional changes in brain activity, while immunohistochemistry for markers of white matter integrity, GABAergic interneurons and glial activation were employed to investigate the most likely related cellular changes. By combining the results from these techniques, we gain a more comprehensive understanding how alterations in the brain can affect cortical activity, ultimately leading to a better understanding of the underlying mechanisms driving mild traumatic brain injury. Importantly, EEG is easy to use, portable, and cost-effective, making it a viable candidate for widespread use in the battlefield and clinical settings. Using a bTBI rodent model, we present the first analysis of electrophysiological, neuronal, axonal and inflammatory pathology in mild bTBI over the acute and chronic phases of injury. While at the acute timepoint, we demonstrate minimal electrophysiological abnormalities, at chronic timepoints our findings reveal a characteristic electrophysiological power spectrum signature of TBI as well as an increase in gamma band functional connectivity. Additionally, we report the first evidence of accompanying changes in cortical regions and layer-specific GABAergic inhibitory interneuron (SST and PV) density.

## Methodology

### Animals

All animal experiments were conducted in compliance with the Home Office licence ([Scientific Procedures] Act 1986 and EU legislation). Male Sprague Dawley rats (∼ 330 g, Charles River, Margate, UK) were housed in individually ventilated cages of four until animals reached 350 g, and subsequently housed in pairs. The facility provided a 12-hour light/dark cycle with temperature and humidity control. Animals were provided with standard rodent chow (Rat and Mouse No.1 Maintenance, Special diets service, UK) and sterile water. All procedures and experiments were conducted during the light hours after a minimum 7-day acclimatisation period.

### Experimental Design

All blast experiments employed the Royal British Legion Centre for Blast Injuries (TRBL-CBIS) shock tube at Imperial College London. No study protocol was registered before the study. For the investigation of the acute effects of injury, 12 animals underwent electrophysiological recordings prior to and after blast exposure in a within-subjects design under continuous terminal anaesthesia. For chronic timepoint investigations, prior to blast exposure, animals were assigned to one of six groups: naïve (*n* = 3), sham-blast (*n* = 27), and blast exposure: 6-hours (*n* = 6), 7-days (*n* = 7), 1month (*n* = 22) and 3-months post-injury *(n* = 26). Prior to blast, these groups were further subdivided into cohorts for either electrophysiological or histological assessment post-injury. No sample size calculations were performed prior to the start of the study, but sample sizes were chosen to be aligned to or larger than those in previous literature.

The histopathological cohort (*n* = 39) was subject to terminal anaesthesia, transcardial perfusion with PBS and 4% paraformaldehyde at different time points post blast (either 6-hours, 7-days, 1month, 3-months post-blast). The electrophysiological cohort (*n* = 50) was subjected to terminal anaesthesia, EEG implantation and recording and euthanasia at different time points post blast (either 1 month or 3 months post-blast). The experimental unit was a single animal. Further details about the shock tube as well as the blast procedure can be found in the supplementary material.

### Blast Exposure

Sprague-Dawley rats were subjected to a mild lateral blast under isoflurane using the TRBL-CBIS shock tube. The animals had their thorax protected during the blast and their heads were not restricted. Each animal was allowed to recover in an oxygen enriched warmed cage, given analgesia for 72 hours post-injury and monitored for adverse effects until the end of experiments. Full details of the blast procedure can be found in the supplementary material.

### Electrophysiological Recordings

#### Surgery

All animals underwent electrophysiological recordings under terminal anaesthesia (urethane, intraperitoneal injection, 2.7 ml/kg, 1.35 g/kg, Sigma-Aldrich).

Once a surgical plane of anaesthesia had been achieved, animals were placed on a stereotaxic apparatus (Kopf Instruments) for surgery. A midline incision was made using a scalpel, followed by clearing of connective tissue via blunt dissection to clear the dorsal surface of the skull. Further details about the preoperative preparations and surgery are included in the supplementary material.

#### Recording Setup

Following surgery, animals were head-fixed to a magnetic arm via a head plate affixed to the skull (caudal of the lambdoid suture) with dental cement and placed inside a shielded soundproofed recording chamber. 32-channel EEG multielectrode arrays (Rat EEG Functional, Neuronexus) were implanted on the dorsal surface of the skull using a saline drop for adhesion, with a reference electrode placed in the neck. The array covered an area from the lambdoid suture to rostral of the coronal suture between the lateral ridges (Fig. 1C). Signals were acquired using Neuronexus headstages, Smartbox^TM^ acquisition board and software (for details please see supplementary material). Spontaneous activity was recorded for 5-minute intervals over the experimental period.

**Figure 1.**
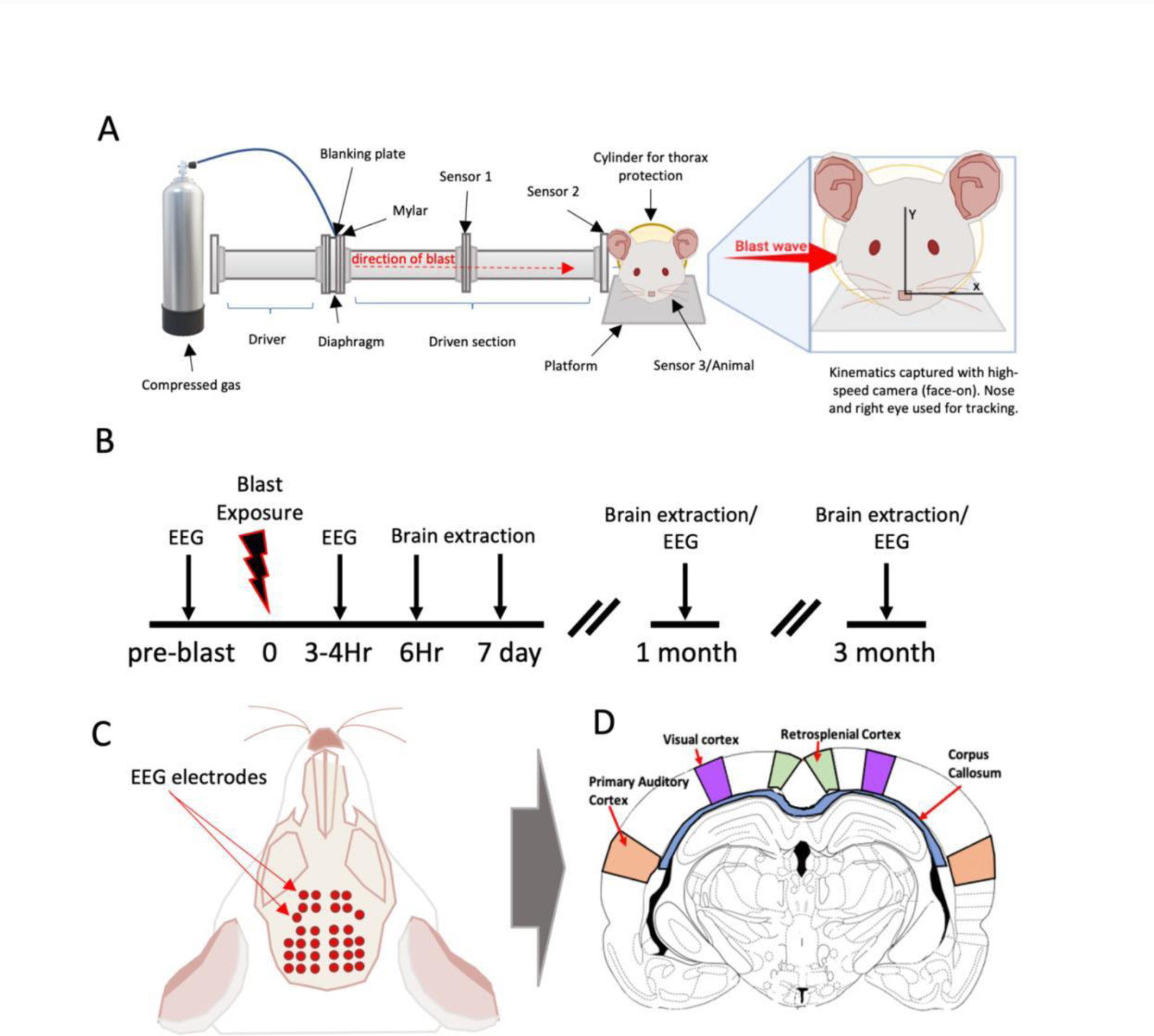
Overview of methodology: A) A schematic of the TRBL CBIS shock tube used for blast exposures. It depicts the compressed gas cylinder that fills 7% of the driver tube using a blanking plate and 250 mm Mylar® diaphragms (which burst when threshold pressure is reached). Sensors 1, 2 and 3 are also visible down the driven shock tube. Sensor 2 is flush with the internal wall of the shock tube whereas sensor 3 was placed during experiments with the animal not present. There is a visualised expansion to the right of the tube schematic which indicates the direction of blast wave. The animal was placed perpendicular to the direction of the blast wave with the right side of the head closest to the shock tube exit. One fiducial point of reference used for tracing the kinematics, the nose, is shown. **B)** Experimental timeline. **C)** Schematic of the electrophysiological set up. Red dots represent the EEG electrodes. Blue lines represent the approximate location of the histological sections relative to the EEG array. **D)** Representative schematic of sections stained. Coronal sections stained for the histological cohort were analysed in these regions of interest. Blue = Corpus callosum, orange = Primary auditory cortex, purple = Primary visual cortex and green = Retrosplenial cortex. Difference between ipsilateral and contralateral hemisphere is shown with a red dotted line. Images were adapted from an atlas (Paxinos and Watson 2007).

### EEG Data Analysis

#### EEG Pre-processing

Data were pre-processed using functions from the EEGLAB toolbox (Delorme and Makeig 2004). Signals were first resampled at 1 kHz and then line noise at 50 Hz and harmonics were suppressed using the CleanLine method version from the PREP toolbox (function *cleanLineNoise*, (BigdelyShamlo et al. 2015)). Data were then scanned for artifacts using the Artifact Subspace Reconstruction algorithm (function *clean_artifacts,* (Mullen et al. 2015)). Segments containing artifacts were discarded from subsequent analysis. Finally, data were decomposed into independent components, using the infomax algorithm (function *pop_runica*, (Bell and Sejnowski 1995)) with default settings. Resulting components were conservatively rejected for physiological artifacts based on their activation timeseries and spectra. Further details on the choice of parameters and preprocessing steps are available in the supplementary material.

#### EEG Analysis of Power

All datasets were ensured to be of equal duration to achieve comparable signal to noise ratios. Power spectral densities were computed for each channel from 1 Hz to 200 Hz using the fast Fourier transform and the Welsch’s method on 2 s consecutive windows with 50% overlap (function *pop_spectopo*), with the resulting spectra normalised to the total power. Normalised power was then obtained for canonical EEG frequency bands (Delta: 1–4 Hz, Theta: 4-8 Hz, Alpha: 8-12 Hz, Beta: 12-25 Hz, Gamma: 25-80 Hz, High-frequency Oscillations: 80-200 Hz) by taking the mean of the values within each band. To obtain global power, spectra were averaged across the electrode array. For the analysis of power at the electrode level, channels rejected during preprocessing (fewer than three channels for any animal) were interpolated using spherical interpolation (function *pop_interp* with default settings; results were identical when an inverse distance method was used). When multiple runs were available per animal, a summary measure of all runs was obtained by averaging over runs.

#### EEG Analysis of Connectivity

Data were epoched into 3 s segments, and the cross-spectral density for each electrode pair computed using Welch’s method with a 1.5 s sliding window with 50% overlap (MATLAB function *cpsd*). These values were used as input to the low-level FieldTrip (Oostenveld et al. 2011) function *ft_connectivity_wpli*, to compute the debiased, weighted phase lag index (dwPLI) (Vinck et al. 2011; Oostenveld et al. 2011), between electrode pairs, a phase synchronisation measure of connectivity robust to volume conduction. Connectivity values were averaged within the frequency bands defined above to assess band-specific differences between groups. To obtain global connectivity, dwPLI values were averaged across all pairs of the electrode array. When multiple runs were available per animal, a summary measure of all runs was obtained by averaging over runs.

#### Immunostaining

Following transcardial perfusion with PBS and 4% paraformaldehyde, brains were postfixed (24 hours), blocked, processed and paraffin-embedded. From paraffinised blocks, sections were cut (7 µm) coronally. To investigate neuronal density, sections were stained with an antibody against NeuN (Millipore), and for specific GABAergic inhibitory interneuron density with Parvalbumin (PV) (Swant) and Somatostatin (SST) (Millipore). To assess changes in the density of glial cells in response to the blast we stained for glial fibrillary acidic protein (GFAP) for astrocytes, and ionised calcium-binding adaptor molecule 1 (IBA1) as a marker for microglia, and visualised glial activation using 3,3′-Diaminobenzidine (DAB) (Vectorlabs). Immunofluorescent staining was conducted with neurofilament light (NFL) to assess axonal damage in the white matter. Luxol fast blue (LFB) was used to analyse the thickness of the corpus callosum (CC) and is a marker of myelin. Details for the immunostaining are found in the supplementary material.

#### Immunostaining Imaging

Full coronal slices were used for light-microscopy (NeuN, PV, SST, IBA1 and GFAP), and imaged at 20x with a slide scanner (Aperio AT2 20x/0.75 NA Plan Apo, Leica Biosystems, Germany). For the immunofluorescent images, half sections including the whole corpus callosum and neocortex were acquired using the tile function on the LSM 780-inverted confocal laser scanning microscope (Carl Zeiss), and Zen software at 10x.

#### Immunostaining Image Analysis

The regions of interest (ROI) analysed were the primary auditory cortex (Au1), primary visual cortex (V1), retrosplenial cortex (RSC), and the CC (Fig. 1D). The densities of NeuN, PV and SST positive neurons were analysed separately for each cortical layer (layer 2/3-6) for each ROI. NeuN analysis densities were manually counted using HALO. PV and SST were quantified using a HALO cytonuclear algorithm. For glial analysis no cortical layers were defined for the Au1, V1 or RSC, and the whole CC was outlined. Astrocytes (GFAP) and microglia (IBA1) percentage area stained were assessed using HALO’s percentage area-stained algorithm. Microglial (IBA1) densities were calculated using HALO’s microglial algorithm. Neurofilament light (NFL) in the CC was measured using HALO’s immunofluorescent intensity algorithm and normalised to an internal control. Luxol fast blue (LFB) CC thickness was measured and averaged per animal then normalised to the Naïve/Sham group and presented as % from sham. For all analysis all groups were compared to the Naïve/Sham group. Further details on immunostaining image analysis are described in the supplementary material.

### Statistical Analysis

Statistical analysis was performed using Matlab^TM^ and GraphPad Prism (9.4.1). Electrophysiological changes were assessed using paired *t*-tests for the acute cohort and 2-way ANOVAs (injury group x post-injury timepoint with interaction term) for the chronic cohort. 2 sample *t*-tests were performed for post-hoc pairwise comparisons in the case of an interaction. Histological changes were assessed using one-way ANOVAs with time post-injury as the factor, followed by post-hoc pairwise comparisons against the control group. Data were presented as ± SEM unless stated otherwise, and the threshold for significance was set at *P* = 0.05. Further details on statistical analyses are provided in the supplementary material.

### Data availability

The data underlying this article cannot be shared publicly due to because of ongoing work. The data will be shared on reasonable request to the corresponding authors.

## Results

### Minimal electrophysiological changes in the acute post-blast period

Analysis of *absolute* (Fig. 2A) and *normalised* (Fig. 2B) global EEG power revealed no differences in resting-state EEG power in any of the bands examined. Analysis of global functional connectivity (Fig. 2C) revealed significantly elevated values at 3-4 hours post-blast compared to baseline (*P* < 0.05; paired *t*-test – 11 degrees of freedom) for the high-frequency oscillations (HFO) band.

**Figure 2.**
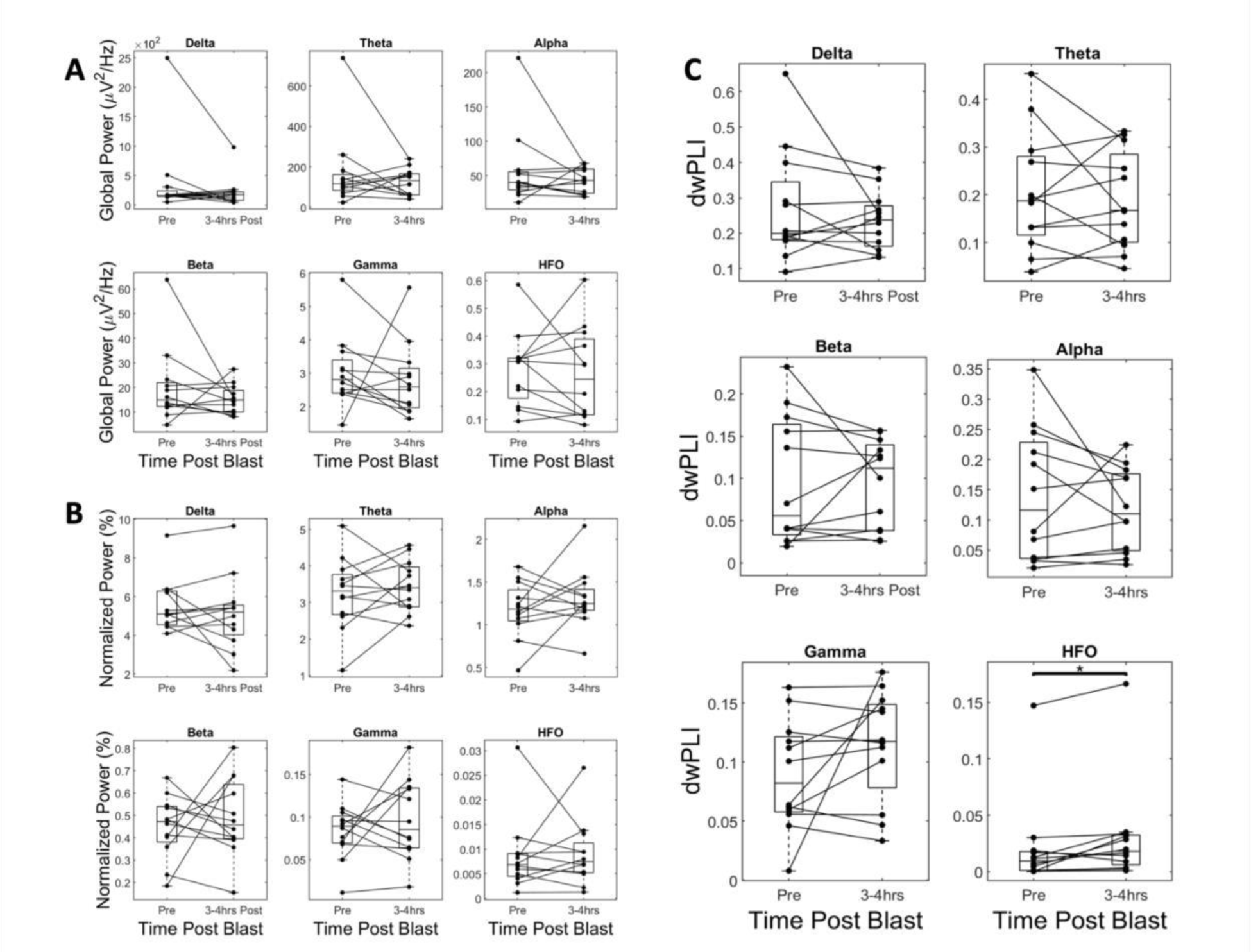
Electrophysiological power and connectivity at the acute post-injury timepoint. (A) Global absolute power across all frequency bands. No significant differences were found for any band. **(B)** Global normalised power across all frequency bands. No significant differences were found for any band. **(C)** Global connectivity across all frequency bands. A significant difference was found for the HFO band. **All panels:** Delta: 1-4 Hz, Theta: 4-8 Hz, Alpha: 8-12 Hz, Beta: 1225 Hz, Gamma: 25-80 Hz, HFO: 80-200Hz. Permutation-based paired *t*-test (Pre-blast Vs 3-4 hours post-blast, 10000 iterations) with Bonferroni correction for multiple comparisons (n = 6 bands) (n = 12 animals).

### EEG power spectrum increases at chronic timepoints due to mild bTBI

Analysis of *absolute* global EEG power revealed a main effect of injury in EEG power (Fig. 3A) (blast vs sham; all cases: two-way ANOVA; *F* (1,46)) with significantly elevated power in the blast groups for most bands examined, (delta: *P* = 0.0138, theta: *P* = 0.0054, alpha: *P* = 0.0222, beta: *P* = 0.0172, gamma: *P* = 0.0006).

**Figure 3.**
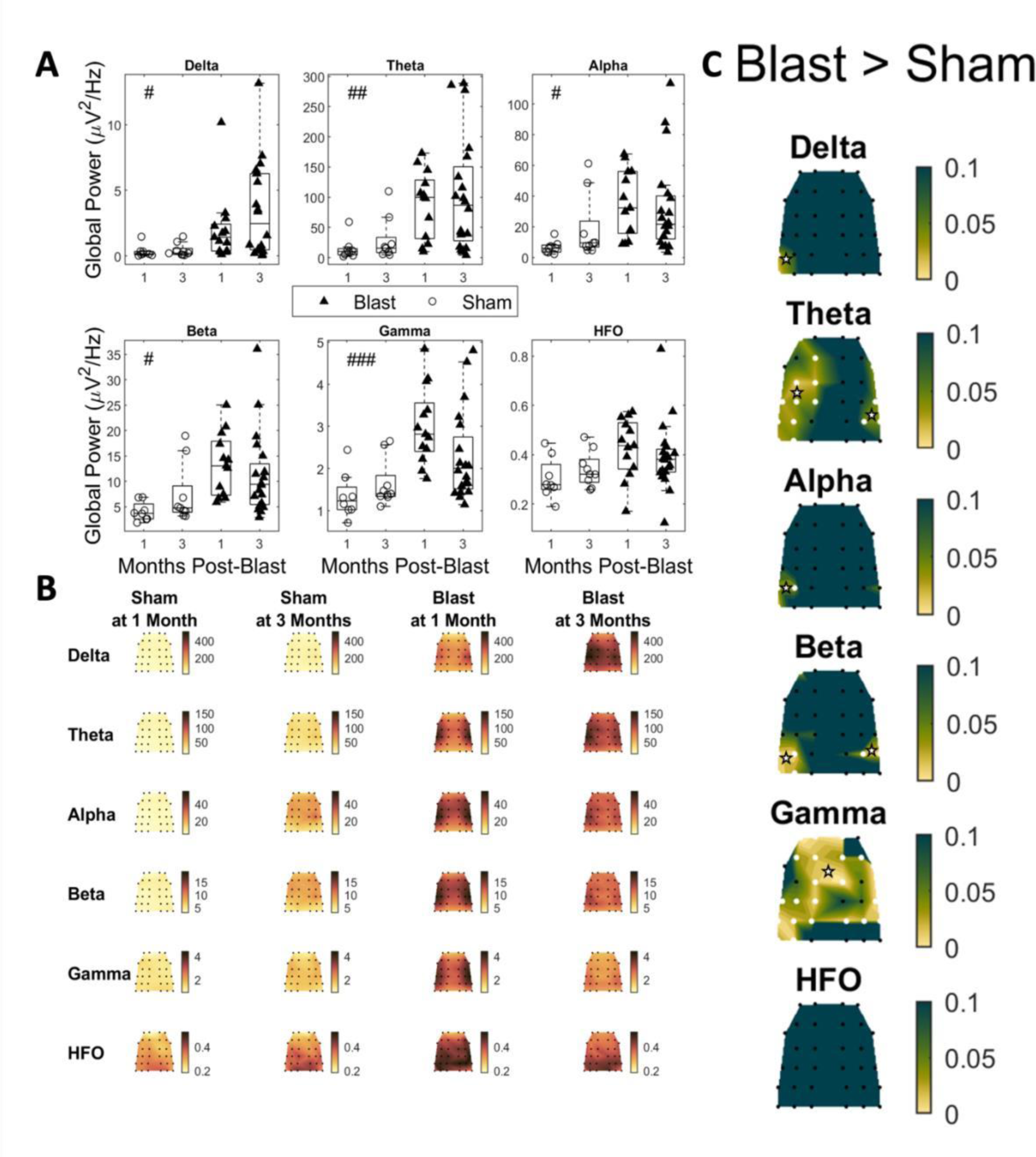
Absolute power in Blast and Sham groups at 1 and -3-months post-injury. (A) Global absolute power for all groups across all frequency bands. Delta: 1-4 Hz, Theta: 4-8 Hz, Alpha: 8-12 Hz, Beta: 12-25 Hz, Gamma: 25-80 Hz, HFO: 80-200 Hz. Permutation-based 2-way ANOVA (Group x Time Post-Blast with interaction, 10000 iterations) with Bonferroni correction for multiple comparisons (n = 6 bands). Significant main effects of group were found for all frequency bands except for the HFO band. Significance values for all tests #: *P*-adj<0.05 main effect of group, ##: *P*-adj<0.01 main effect of group, ###: *P*-adj<0.001 main effect of group, **(B)** Topoplots depicting mean group absolute power for all groups and frequency bands, across the multielectrode array. **(C)** Topoplots depicting *P*-values for the main effect of group (permutationbased 2-way ANOVA, 10000 iterations), Bonferroni corrected for multiple comparisons (n = 32 channels) for all bands. Channels for an adjusted *P*-value below 0.05 are marked with white stars (blast 1-month: n = 13, blast 3-month: n = 20, sham 1-month n= 8, sham 3 -month n = 9).

To further localise the effect, we tested the main effect of injury (all cases: two-way ANOVA; *F* (1,46)) at the electrode level for all the frequency bands of interest (Fig. 3 B,C). An increase in power in the delta band was observed for the blast group in the temporal region of the side contralateral (left) to the blast, as well as in the theta band in temporal, medial and frontal electrodes of the left hemisphere, and in the temporal electrodes of the right hemisphere. Power increases in the left temporal region were also observed in the alpha band for the blast group. In the beta band, the blast group had increased power along the left and right temporal regions. Finally, in the gamma band, the blast group displayed increased power in medial, temporal and frontal regions in both hemispheres. No spatially localised effect of injury was detected for the HFO band.

Analysis of global *normalised* EEG power (Fig. 4A) revealed a significant main effect of injury (blast vs sham; all cases: two-way ANOVA; *F* (1,46)) in most bands examined, driven by significantly elevated power after blast in the low frequency (delta: *P* = 0.0132, theta: *P* = 0.0498) and decreased power for the high frequency bands (beta: *P* = 0.015, gamma: *P* = 0.003, HFO: *P* < 0.001). In the alpha band, a significant interaction was detected (*F* (1,46), *P* = 0.0246), and pairwise comparisons (unequal variances *t*-test; degrees of freedom: *ν* = 18.9076, calculated using Satterthwaite’s approximation) revealed a significant decrease in power for the blast group at 3months compared to the sham group at 3-months (*P* = 0.0105).

**Figure 4.**
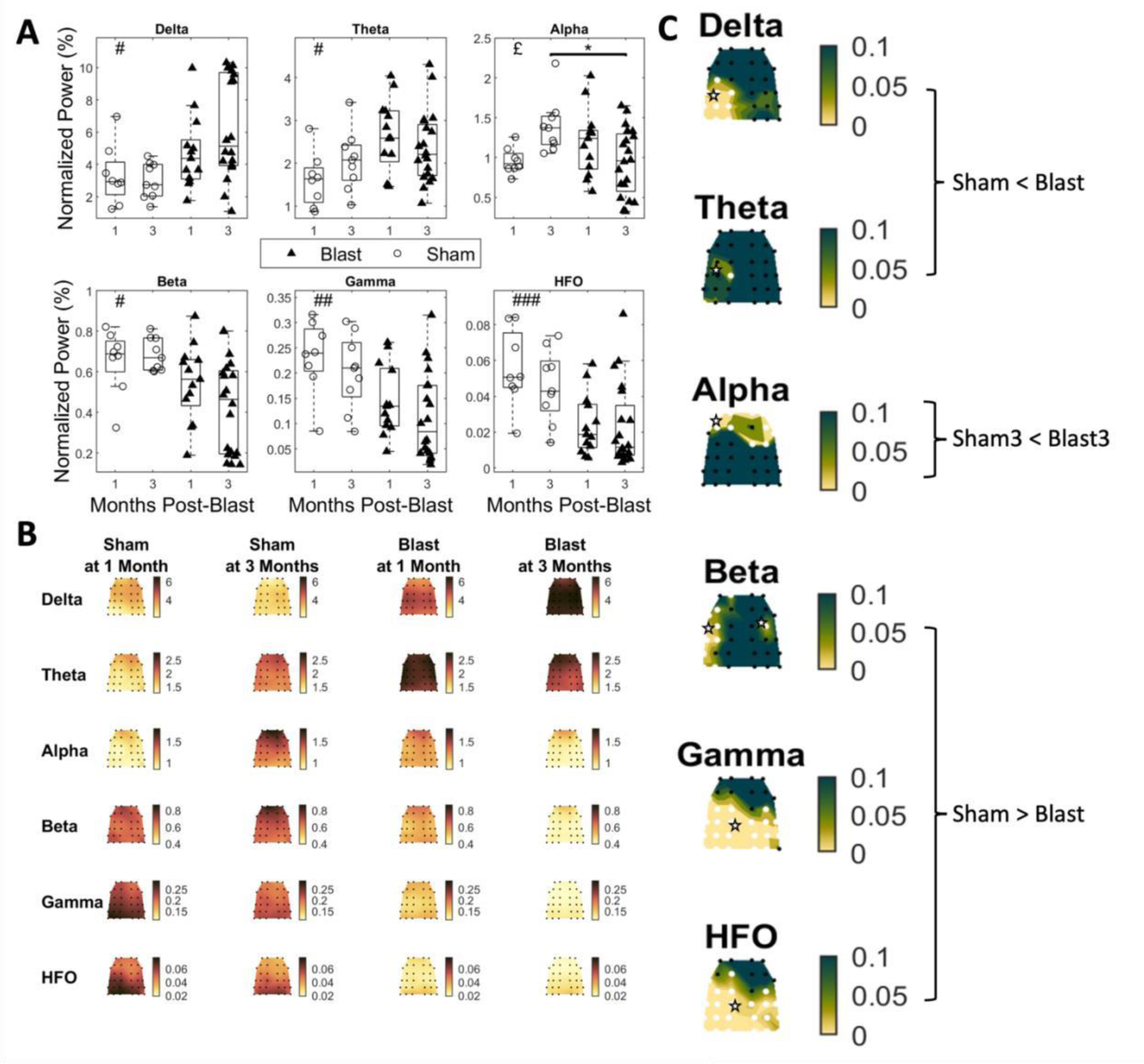
Normalised power in Blast and Sham groups at 1 and 3-months post-injury. (A) Global normalised power for all groups across all frequency bands. Delta: 1-4 Hz, Theta: 4-8 Hz, Alpha: 8-12 Hz, Beta: 12-25 Hz, Gamma: 25-80 Hz, HFO: 80-200 Hz. Permutation-based 2-way ANOVA (Group x Time Post-Blast with interaction, 10000 iterations) with Bonferroni correction for multiple comparisons (*n* = 6 bands). Significant main effects of group were found for all frequency bands except for the alpha band. Significant interaction in the alpha band was followed by pairwise permutation-based *t*-tests (10000 permutations) between the groups of interest with Bonferroni correction for multiple comparisons (n = 3 pairs). Significance values for all tests #: *P*adj<0.05 main effect of group, ##: *P*-adj<0.01 main effect of group, ###: *P*-adj<0.001 main effect of group, £: *P*-adj<0.05 interaction effect : *P*-adj<0.05 two sample *t*-test. **(B)** Topoplots depicting mean group normalised power for all groups and frequency bands, across the multielectrode array. **(C)** Topoplots depicting *P*-values for the main effect of group (permutation-based 2-way ANOVA, 10000 iterations), Bonferroni corrected for multiple comparisons (*n* = 32 channels) for the Delta, Theta, Beta, Gamma and HFO bands. The topoplot for the alpha band depicts *P*-values for the two- sample t-test between sham at 3-months and blast at 3-months. Channels for which the adjusted *P*- value is below a threshold of 0.05 are marked with white stars (blast 1-month: n = 13, blast 3- month: n = 20, sham 1-month: n = 8, sham 3-month: n = 9).

To further localise the effect, we performed testing for the main effect of injury group (all cases: two-way ANOVA; *F* (1,46)) at the electrode level for all the frequency bands of interest and testing for the difference between the sham and blast groups at 3-months for the alpha band (unequal variances *t*-test; degrees of freedom: *ν* = 18.9076, calculated using Satterthwaite’s approximation) (Fig. 4 B,C). For the delta band, normalised power was increased (*P <* 0.05) for the blast group in a region covering the temporal and medial region of the left hemisphere. Normalised power was also elevated for the theta band for the blast group in the left medial region. For the alpha band, the blast group at 3-months had reduced normalised power in frontal regions compared to the sham group at 3-months. In the beta band, the blast group had reduced normalised power in locations along the left temporal and parietal regions. Finally, for the gamma and HFO bands, the pattern was similar, with the blast group having widespread decreased normalised power in posterior regions, spanning medial and temporal regions for both bands and an additional left frontal region for the HFO band.

### Mild blast injury increases Functional Connectivity in the Gamma Band at chronic timepoints

Analysis of global connectivity for all bands of interest (Fig. 5A, all cases: two-way ANOVA; *F* (1,46)), revealed significantly elevated values for the blast group specific to the gamma band (injury group effect *P* = 0.0108). After pooling the 1-month and 3-months datasets for both injury groups and conducting further analysis at the edge level for the gamma networks, a robust network (one-sided independent samples *t*-test; *t* (49), *P* < 0.05) of increased connectivity in the blast group was obtained (Fig. 5B). This network included both interhemispheric and intrahemispheric connections spanning frontal, central, temporal, and medial regions.

**Figure 5.**
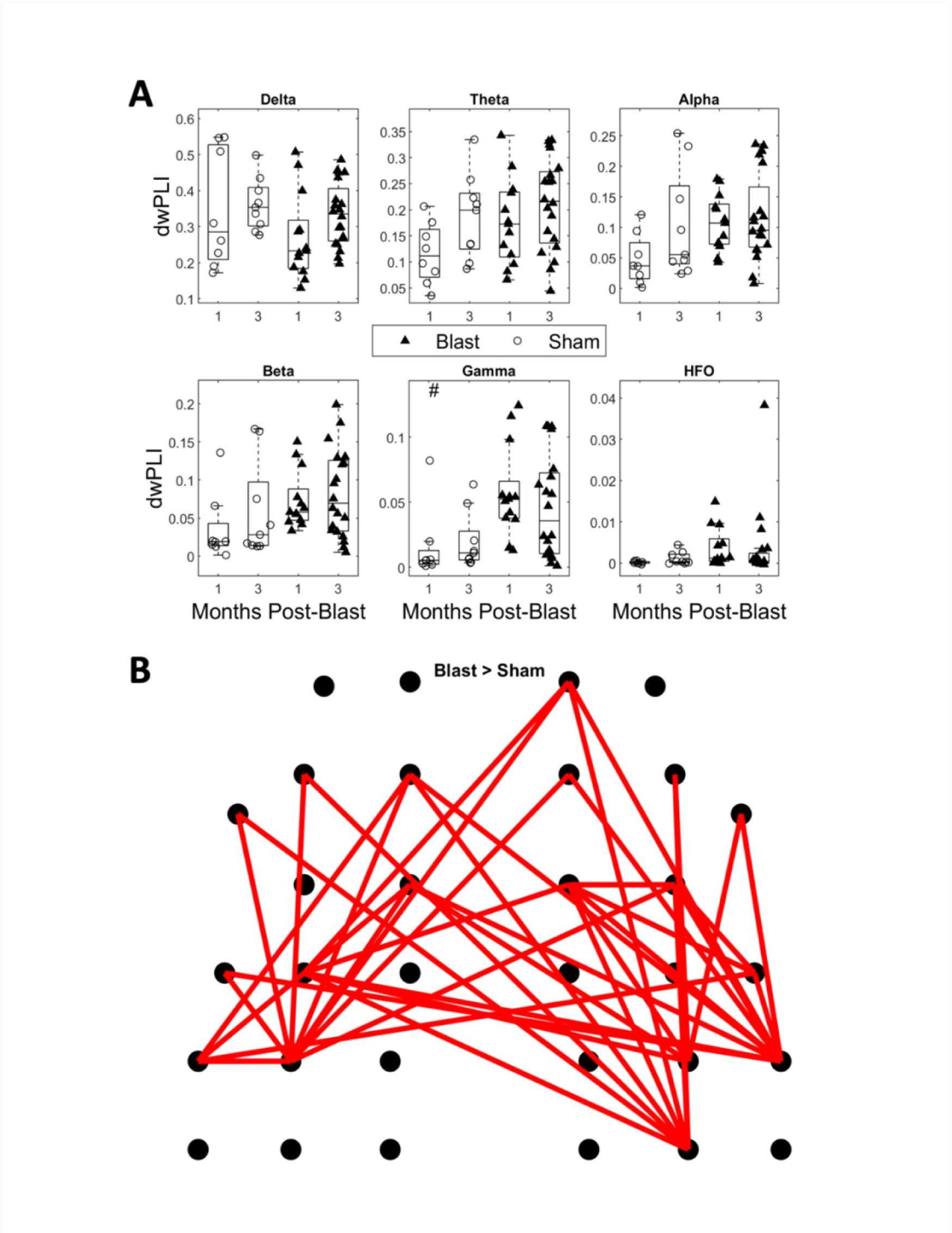
Phase-based functional connectivity (dwPLI) in Blast and Sham groups at 1 and 3months post-injury. (A) Global connectivity for all groups. Permutation-based 2-way ANOVA (Group x Time Post-Blast with interaction, 10000 iterations) with Bonferroni correction for multiple comparisons (n = 6 bands). A significant main effect of group was found only for the gamma band. Significance values for all tests : *P*-adj<0.05. **(B)** Topoplot depicting a significantly hyperconnected network for the blast group. Independent-samples t-test (blast vs sham) based on 50000 permutations with an FDR-adjusted threshold of *P* = 0.05, using the Network-based statistic toolbox (blast 1-month: n = 13, blast 3-month: n = 20, sham 1-month: n = 8, sham 3-month: n = 9).

### Mild Blast Exposure Decreases layer and region-specific PV and SST Interneuronal density at 1 month and 3 months post-blast

We performed NeuN staining as a marker of neurons to determine whether electrophysiological changes were associated with alterations in gross scale cortical neuronal density. Our results revealed no effects on overall neuronal number in the ROIs (Fig. 6).

**Figure 6:**
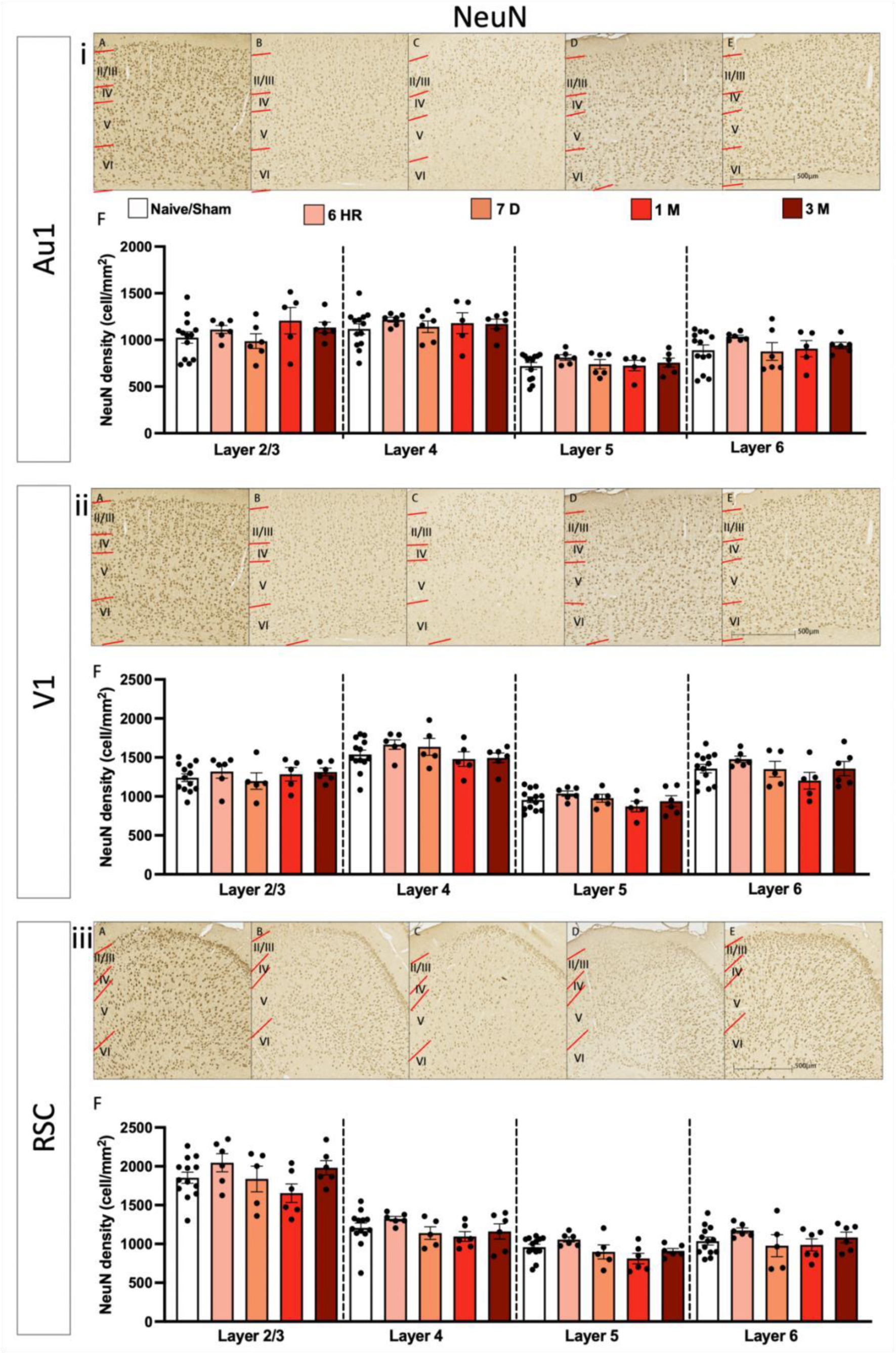
NeuN density changes in the primary auditory cortex, the primary visual cortex and the retrosplenial cortex post-blast traumatic brain injury. Density of NeuN positive cells following single blast traumatic brain injury in the primary auditory cortex for **(iA-iB)** Naïve/Sham, **(iC-iD)** 6-hours post-injury, **(iE-iF)** 7-days post -injury, **(iG-iH)** 1-month postinjury, **(iI-J)** 3-months post-injury. **(iA/iC/iE/iG/iI)** Micrograph of all cortical layers in the primary auditory cortex. **(iB/iD/iF/iH/iJ)** higher magnification micrographs of 5th and 6th layer of the cortex. **(iK-iN)**. NeuN density in the primary visual cortex for **(iiA-iiB)** Naïve/Sham, **(iiCiiiD)** 6- hours post-injury, **(iiEi-iiF)** 7-days post -injury, **(iiG-iiH)** 1-month post-injury, **(iiI-iiJ)** 3months post-injury. **(iiA/iiC/iiE/iiG/iiI)** Micrograph of in the 5th and 6th layer of the primary visual cortex **(iiiK-N)**. **(iiiB/iiiD/iiiF/iiiH/iiiJ)** higher magnification micrographs. NeuN density in the retrosplenial cortex for **(iiiA-B)** Naïve/Sham, **(iiiC-iiiD)** 6-hours post-injury, **(iiiE-F)** 7-days post -injury, **(iiiG-H)** 1-month post-injury, **(iiiI-iiiJ)** 3-months post-injury. **(iiA/iiiC/iiiE/iiiG/iiiI)** Micrograph of all cortical layers in the retrosplenial cortex. **(iiiB/iiiD/iiiF/iiiH/iiiJ)** higher magnification micrographs **(iiiK-iiiN)** quantification of interneuron cell density using one-way ANOVA, corrected using BKY FDR. All data presented mean ± SEM. \**P*<0.05, \*\**P*<0.01, \*\*\**P*<0.001 and \*\*\*\**P*<0.0001 (Au1: Naïve/Sham: n = 13, 6 Hr: n = 6, 7-day: n = 6, 1-month: n = 5, 3-months: n = 6. V1: Naïve/Sham: n = 13, 6 Hr: n = 6, 7-day: n = 5, 1-month: n = 5, 3-months: n = 6. RSC: Naïve/Sham: n = 13, 6 Hr: n = 6, 7-day: n = 5, 1-month: n = 5, 3-months: n = 6).

We then investigated whether specific neuronal types might be affected by blast exposure. As mentioned, PV and SST interneurons have been shown to be preferentially damaged by TBI and were therefore selected for histological analysis. GABAergic PV INs were analysed in three cortical regions and only the 5^th^ and 6^th^ layer of the Au1 was found to be significantly affected (layer 5: *F* (4, 33) = 5.291, *P* = 0.0021, BKY FDR adjusted-*P* value: *q* = 0.0084, layer 6: *F* (4, 33) = 4.670, *P* = 0.0043, *q* = 0.0172: Fig. 7i). Further analysis revealed a significant decrease in PV INs density at 7-days and 1-month post-injury for both the 5^th^ (7-days: *P* = 0.0084, 1-month: *P* = 0.0006, a reduction of 29.53% for 7-days and 45.73% for 1-month post-injury relative to control) and 6^th^ cortical layers (7-days: *P* = 0.0041, 1-month: *P* = 0.0005, a reduction of 33.88% and 47.63% respectively). There were no significant changes in PV IN density in any cortical layer for both the V1 (Fig. 7ii) and the RSC (Fig. 7iii), *P* and *q* >0.05.

**Figure 7:**
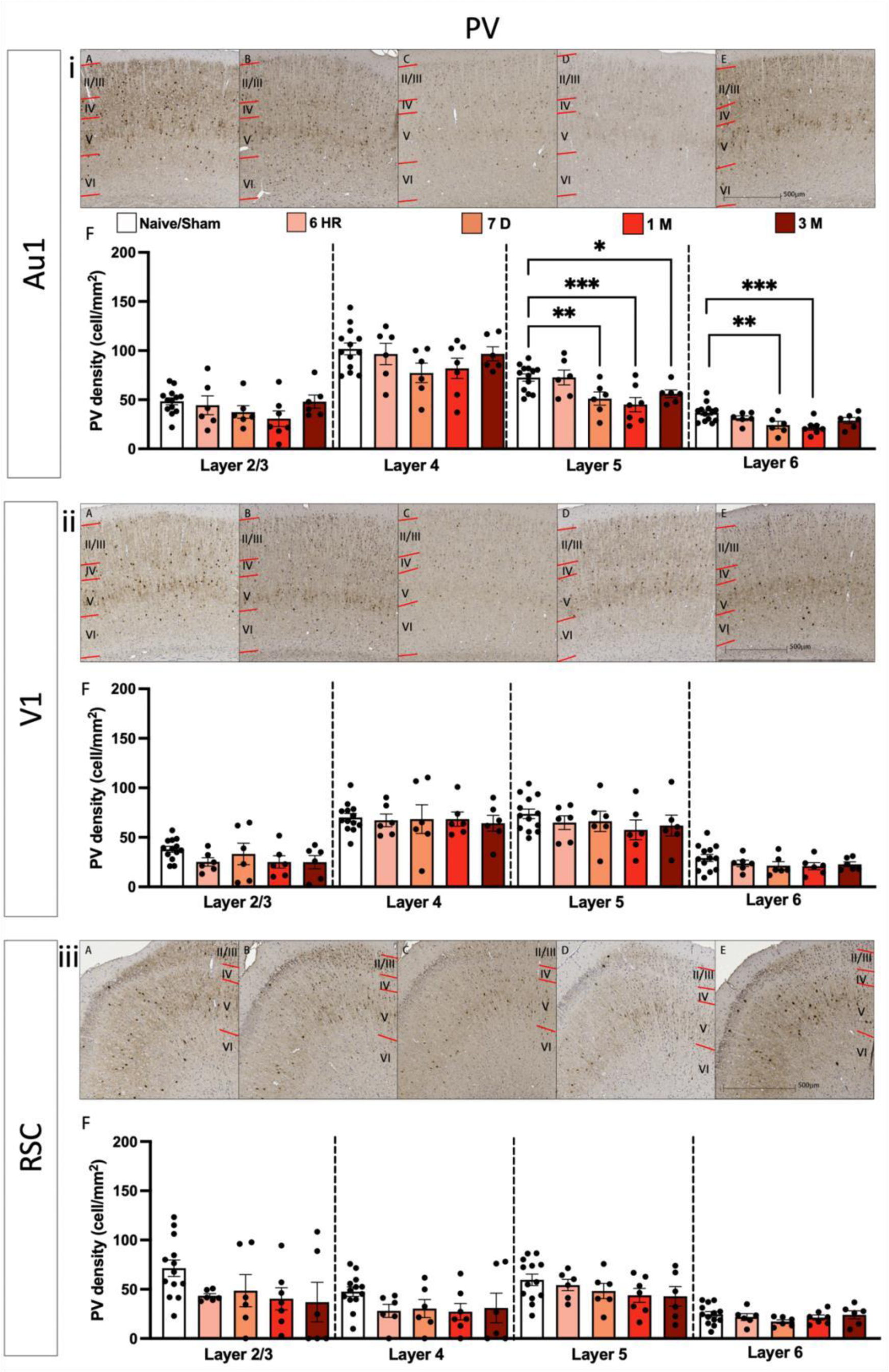
Parvalbumin density changes in the primary auditory cortex, the primary visual cortex and the retrosplenial cortex following blast TBI. Density of Parvalbumin positive cells following single blast traumatic brain injury in the primary auditory cortex **for (iA-B)** Naïve/Sham, **(1C-D)** 6-hours post-injury, **(iE-F)** 7-days post -injury, **(iG-H)** 1 month post-injury, **(iI-J)** 3months post-injury. **(iA/iC/iE/iG/iI)** Micrograph of all cortical layers in the primary auditory cortex. **(iB/iD/iF/iH/iJ)** higher magnification micrographs of 5th and 6th layer of the cortex. **(iKN)**. Parvalbumin density in the primary visual cortex primary for **(iiA-B)** Naïve/Sham, **(iiC-D)** 6hours post-injury, **(iiE-F)** 7-days post -injury, **(iiG-H)** 1-month post-injury, **(iiI-J)** 3-months postinjury. **(iiA/iiC/iiE/iiG/iiI)** Micrograph of all cortical layers in the primary visual cortex **(iiiK-N)**. **(iiiB/iiiD/iiiF/iiiH/iiiJ)** higher magnification micrographs. Parvalbumin density in the retrosplenial cortex for **(iiiA-B)** Naïve/Sham, **(iiiC-D)** 6-hours post-injury, **(iiiE-F)** 7-days post injury, **(iiiG-H)** 1-month post-injury, **(iiI-J)** 3-months post-injuryi. **(iiA/iiiC/iiiE/iiiG/iiiI)** Micrograph of all cortical layers in the retrosplenial cortex. **(iiiB/iiiD/iiiF/iiiH/iiiJ)** higher magnification micrographs **(iiiK-N)** quantification of interneuron cell density using one-way ANOVA, corrected using BKY FDR. All data presented mean ± SEM. \**P*<0.05, \*\**P*<0.01, \*\*\**P*<0.001 and \*\*\*\**P*<0.0001 (Au1: Naïve/Sham: n = 13, 6 Hr: n = 6, 7-day: n = 6, 1-month: n = 7, 3-months: n = 6. V1: Naïve/Sham: n = 13, 6 Hr: n = 6, 7-day: n = 6, 1-month: n = 7, 3-months: n = 6. RSC: Naïve/Sham: n = 13, 6 Hr: n = 6, 7-day: n = 6, 1-month: n = 6, 3-months: n = 6).

The density of SST positive cells in the 5^th^ layer of the neocortex was found to be significantly reduced at specific time points post-injury. One-way ANOVAs, corrected using previously mentioned multiple-comparison correction method, revealed significant differences in the 5^th^ layer in the Au1 (*F* (4, 34) = 4.140, *P* = 0.0077, *q =* 0.024), V1 (*F* (4, 34) = 4.241 *P* = 0.0068, *q =* 0.0214) and RSC (*F* (4, 34) = 3.069, *P* =0.0292, *q* = 0.0460). Further analysis showed that densities decreased from 7-days post-injury relative to Naïve/Shams in the Au1 (Fig. 8i) (7-days: *P* = 0.0029, 1-month: *P* = 0.0147, 3-months: *P* = 0.0085, and amounted to a reduction of 35.52% for 7-days, 25.89% for 1-month, and 30.91% for 3-months). Similar changes were observed in the V1 with density decreases from 7-days post-injury (Fig. 8ii) (7-days: *P* = 0.0025 1-month: *P* = 0.0029, 3- months: *P* = 0.0124 which constitutes a density reduction of 40.34%, 36.09% and 32.55% respectively). A post-hoc test revealed a significantly lower cell density at 3-months post-injury in the 5^th^ layer of the RSC compared to control (*P =* 0.0201, a 29.47% reduction relative to Naïve/Sham). A significant decrease in cell density was found in 2/3^rd^ layer of the RSC (*F* (4, 34) = 5.010 *P* = 0.0028, *q* = 0.0088) with 1-month and 3-months post-injury driving the change (1month: *P* = 0.0022, 3-months: *P* = 0.0252, a density reduction of 43.35% and 33.61% respectively relative to Naïve/Sham) (Fig. 8iii). No significant differences were found in other layers of the other cortical ROIs (*P* and *q* > 0.05).

**Figure 8:**
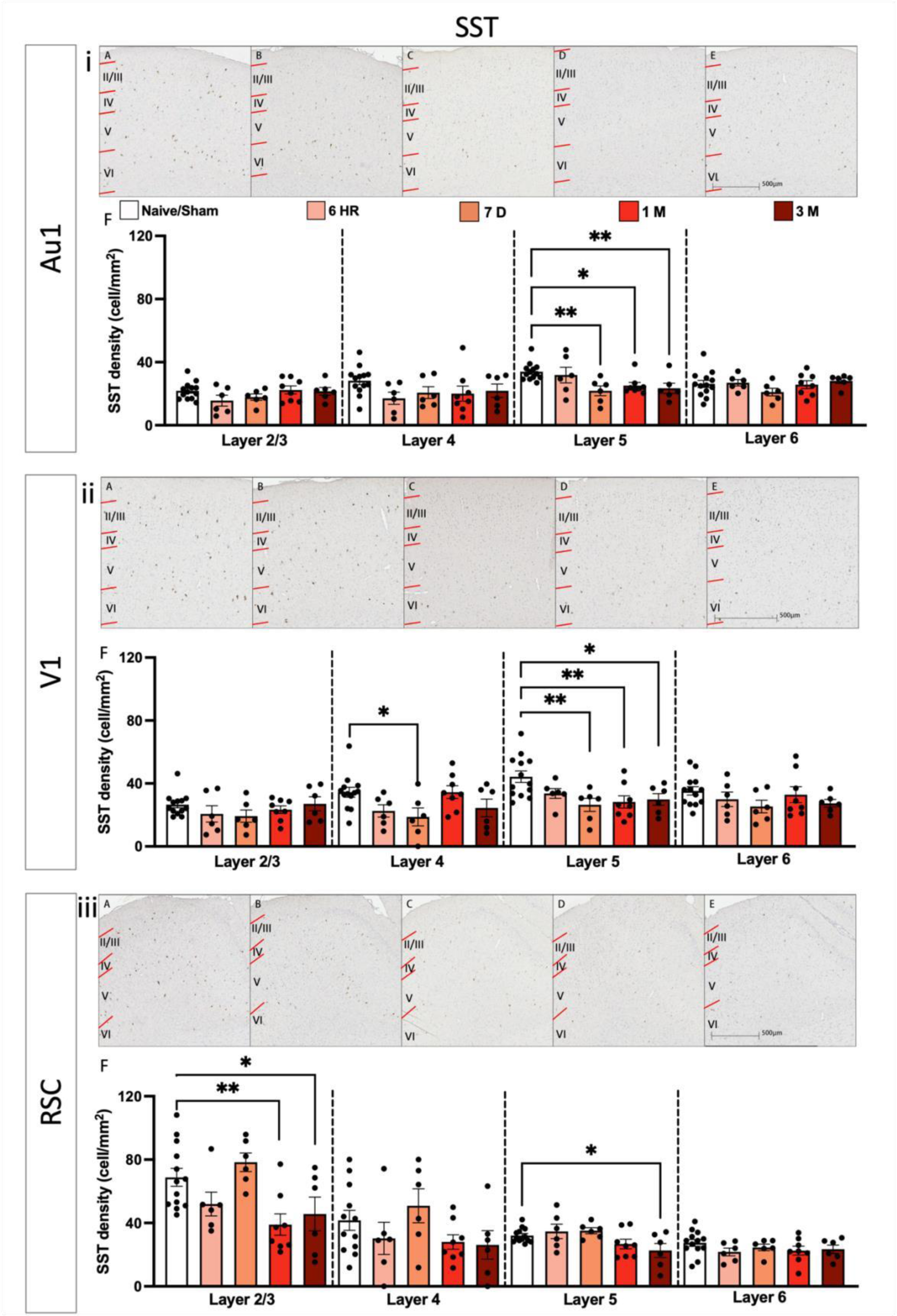
**Somatostatin density changes in the primary auditory cortex, the primary visual cortex and the retrosplenial cortex post-blast traumatic brain injury**. Density of Somatostatin positive cells following single blast traumatic brain injury in the primary auditory cortex at **(iA-B)** Naïve/Sham, **(1C-D)** 6-hours post-injury, **(iE-F)** 7-days post -injury, **(iG-H)** 1-month post-injury, **(iI-J)** 3-months post-injury. **(iA/iC/iE/iG/iI)** Micrograph of all cortical layers in the primary auditory cortex. **(iB/iD/iF/iH/iJ)** higher magnification micrographs of 5th and 6th layer of the cortex. **(iK-N)**. Somatostatin density in the primary visual cortex for **(iiA-B)** Naïve/Sham, **(iiC-D)** 6-hours post-injury, **(iiE-F)** 7-days post -injury, **(iiG-H)** 1-month post-injury, **(iiI-J)** 3-months post-injury. **(iiA/iiC/iiE/iiG/iiI)** Micrograph of in the 5th and 6th layer of the primary visual cortex **(iiiK-N)**. **(iiiB/iiiD/iiiF/iiiH/iiiJ)** higher magnification micrographs. Somatostatin density in the retrosplenial cortex for **(iiiA-B)** Naïve/Sham, **(iiiC-D)** 6-hours post-injury, **(iiiE-F)** 7-days post injury, **(iiiG-H)** 1-month post-injury, **(iiI-J)** 3-months post-injury. **(iiiA/iiiC/iiiE/iiiG/iiiI)** Micrograph of all cortical layers in the retrosplenial cortex. **(iiiB/iiiD/iiiF/iiiH/iiiJ)** higher magnification micrographs **(iiiK-N)** quantification of interneuron cell density using one-way ANOVA, corrected using BKY FDR. All data presented mean ± SEM. \**P*<0.05, \*\**P*<0.01, \*\*\**P*<0.001 and \*\*\*\**P*<0.0001 (Au1: Naïve/Sham: n = 13, 6 Hr: n = 6, 7-day: n = 6, 1-month: n = 8, 3-months: n = 6. V1: Naïve/Sham: n = 13, 6 Hr: n = 6, 7-day: n = 6, 1-month: n = 8, 3-months: n = 6. RSC: Naïve/Sham: n = 13, 6 Hr: n = 6, 7-day: n = 6, 1-month: n = 8, 3-months: n = 6).

### Mild Blast Exposure Causes Microglial changes at Acute and Chronic Timepoints

To examine whether an inflammatory response accompanied the PV and SST loss after blast exposure, glia densities (IBA1- and GFAP-positive cells) were assessed in the same cortical areas. Significant increases in microglia density were observed in the Au1 (*F* (4, 27) = 4.392, *P* = 0.0073) and RSC (*F* (4, 27) = 5.504 *P* = 0.0023), but not the V1 or the CC (*P >*0.05). Further analysis revealed an increase of 34% at 6-hours (*P* = 0.0004) and 18% at 1-month post-injury (*P* = 0.0414) for the Au1 driving the change in IBA1 density, while for the RSC significant differences were found at 6-hour (40% increase *P* = 0.0001), 7-days (25% increase, *P =* 0.0207) and 1-month (20% increase *P* = 0.0288) compared with Naïve/Sham animals. Despite the non-significant main effect for the other regions, there was a trend of increased IBA1 density at 6-hours in the CC and RSC (Supplementary Fig. 2).

Results for percentage IBA1 area covered was found to be significant in the AU1, (F (4,26 = 2.983), *P* = 0.0417), V1 (F (4,26 = 3266), *P* = 0.0269), and CC (F (4,26 = 5.324, *P* = 0.0029), with only borderline differences in the RSC (F (4, 26) = 2.374, *P*=0.0782). Post-hoc analysis revealed that there was an increase in the IBA1 percentage area stained at chronic time points post-injury in all brain areas investigated, as shown in Supplementary Fig. 3 (Au1: area increase of 96.82%, *P* = 0.0326, V1: 106.43% increase, *P* = 0.0250, CC: 266.38% increase, *P* = 0.0006) and the RSC followed the same increasing trend at this chronic timepoint (Supplementary Fig.3).

GFAP area covered as measurement of reactive astrogliosis showed no significant increases in the Au1, V1, RSC and CC (*P* > 0.05) (Supplementary Fig. 4).

### Mild Blast injury affects Corpus Callosum integrity without white matter thinning

NFL is a fundamental neuronal scaffolding protein and has been shown to indicate axonal damage and neurodegeneration in TBI and other neurodegenerative diseases (Graham et al. 2021).

A significant increase in NFL staining intensity was observed in the CC at 1-month post-injury (*P* = 0.0008, a 253% increase) when compared to the control group (Supplementary Fig. 5).

Luxol fast blue was also used to measure for the presence of post-injury CC thinning. A one-way ANOVA showed no differences between any time post-injury and the Naïve/Sham group (Supplementary Fig. 6).

## Discussion

This study investigates the potential of a clinically translationally technique, high-density EEG, as a diagnostic tool in an animal model of mild bTBI. This is the first study to combine cellular pathology, multi-electrode high-density EEG recordings at the whole brain level, and changes in SST interneuron cell density over multiple timepoints from the acute to chronic phase in a model of mild blast-induced TBI.

Our results demonstrate that a single mild blast injury can result in chronic global increases in power and increased gamma band functional connectivity. These electrophysiological signatures of mild bTBI were accompanied by significant decreases across all later timepoints in key inhibitory neural population densities, specifically SST interneuron density in layer 5 across all cortical ROIs, and PV interneuron density in layer 5 of the primary auditory cortex. Additionally, we have shown a mild inflammatory response to blast TBI, as well as indications of chronic white matter disruption.

We only observed minimal effects of blast on electrophysiological metrics of brain activity at the acute timepoint. The time of recording (3-4 hours post injury) is consistent with the literature that supports a return of the EEG signal to baseline 1-hour after injury (see (Ianof and Anghinah 2017) for a review of the time course of EEG changes after TBI). This suggests that our observed chronic changes could be a result of mechanisms acting at longer timescales, likely related to secondary injury mechanisms after TBI. This also suggests the availability of a therapeutic window at this stage. We did however observe significantly elevated functional connectivity in the HFO band. Pathologic activity in this band has been hypothesised as an early precursor to epilepsy (Bragin et al. 2004; Reid et al. 2016) in the pathological cascade driven by hyperexcitability (Golub and Reddy 2022). In our model, this could be driven by an excessive release of excitatory neurotransmitters and accumulation of calcium, both well-documented stages of the post-injury cascade (Guerriero, Giza, and Rotenberg 2015). Such a mechanism is also a putative driver of the increase in inflammatory markers at 6-hours after injury (Simon et al. 2017) that we observe here (Supplementary Fig. 2), with the potential to establish a positive feedback loop of hyperexcitability and neuroinflammation (Sharma et al. 2019; Webster et al. 2017).

Hyperexcitability and an imbalance in excitation and inhibition is a persistent sequela of TBI and it has been proposed that extra-cranial electrophysiology can be used as a tool to monitor these changes in neuronal functionality (Carron, Alwis, and Rajan 2016). Most human studies quantify EEG in terms of *relative* (normalised to total subject power) power changes (see (Mallas et al. 2022)). This is to account for the large degree of individual variability found in total power in human EEG studies (Wang, Ethridge, et al. 2017). This approach, while understandable, will miss important patterns of effects such as a uniform increase in power across all frequencies. Due to the reduction of individual variability available in a preclinical laboratory environment, our model was able to add increased explanatory power to clinically relevant findings by including absolute measures of change. Our analysis in terms of relative power (Fig. 4) yielded similar results to human studies (Mallas et al. 2022), where a relative increase in low frequencies and a decrease in the higher frequencies was observed. However, we were also able to show that this was accompanied by a broadband increase in absolute power (Fig. 3). This suggests the presence of hyperexcitability in our mild bTBI model in the chronic post-injury phase, as similar patterns of increased power have been observed in a knockout model of Fragile-X syndrome (Jonak et al. 2020), a condition linked to circuit hyperexcitability (Contractor, Klyachko, and Portera-Cailliau 2015). Crucially, the two analysis approaches converge in terms of increases in low frequency power, which is a hallmark signature of TBI (Huang et al. 2014; Mallas et al. 2022). Additionally, we have identified an absolute measure of power change as a possible EEG signature of neural hyperexcitability.

Several human studies have established an increase in power in lower frequencies as a hallmark electrophysiological signature of TBI (Robb Swan et al. 2015; Huang et al. 2014; Modarres et al. 2016; Davenport et al. 2022; Safar et al. 2021). This has been linked to cognitive impairment (Mallas et al. 2022) and structural disconnection due to axonal injury (Huang et al. 2014). In our results, we show this same typical electrophysiological signature (Fig. 3, Fig. 4), accompanied by increased neurofilament staining intensity, indicating structural integrity damage in the connecting white matter (Fig. 8G). This lends support to this structural damage being part of the mechanistic substrate underlying the electrophysiological changes.

In line with recent efforts to boost the translational value in preclinical models of neurological disease and align them with modern human electrophysiology studies (Jonak et al. 2018), we used a high-density EEG array to record brain activity. This allowed us to explore electrophysiological correlates of blast TBI both on a global and local scale. With this, we were able to identify regionalised effects of absolute power for the delta, theta, alpha and beta bands in posterior temporal regions, while for the gamma band the effect was more widespread across the array. For relative power, effects for the delta, theta, beta and gamma bands were localised in a similar pattern. Alpha band effects were concentrated in frontal areas, with HFO band effects widespread across central and posterior regions. This pattern of results lends an added degree of spatial sensitivity to our analysis, allowing detection of region-specific signatures of injury and their putative asymmetries across cortex. Future studies could explore the causes of these changes by studying the injury biomechanics through computational modelling (Donat et al. 2021), and take this spatial heterogeneity into account when designing therapeutic interventions.

Functional connectivity studies in TBI show inconsistent findings (Barr et al. 2012; Lewine et al. 2019; Mallas et al. 2022) and this extends to the small number of studies that have investigated functional connectivity changes in blast cohorts. Sponheim et al. (Sponheim et al. 2011) and Wang et al (Wang, Costanzo, et al. 2017) both found reduced FC due to blast but were inconsistent regarding the frequency bands where this reduction was found. In contrast to other studies, and in line with our findings, Huang et al (Huang et al. 2017) reported an *increase* in FC in a human study and found this in the gamma and beta frequency bands. One suggested reason for this seemingly conflicting finding was that subjects in the Huang study were more likely to have suffered a *milder* form of injury and this may have triggered a different pathological path to that found in the more severe injury types typically studied. Indeed, our similar finding in this animal model of *mild* bTBI lends support to this hypothesis.

*Functional hyperconnectivity* has been observed to continue from the acute (24 hours; Manning (Manning et al. 2017)) to the chronic phase (2.5 years on average but up to 6.6 years; Sharp (Sharp et al. 2011)) post-TBI injury, suggesting a protracted elevated shift in baseline connectivity. This is consistent with our results showing hyperconnectivity up to 3-months post injury in rats, roughly corresponding to 1-9 years post-injury in ‘human time’ (Agoston 2017). An increase in functional connectivity as a response to TBI has been correlated to cognitive performance with evidence in favour of it being an adaptive phenomenon (Sharp et al. 2011; Morelli et al. 2021), with the exact relationship to cognition likely depending on injury type, timepoint of assessment, methods, region/network, and cognitive demand (for example, see the mixed correlations with behavioural measures in Huang ^(Huang^ ^et^ ^al. 2017)^). An intriguing interpretation suggests that increased transmission across regions might be a compensatory effect associated with an increase in the efficiency of information exchange across the cortex (Fagerholm et al. 2016). This increase in connectivity could be an attempt to maintain adequate signal to noise ratios in response to the injury (Hillary and Grafman 2017). However, it is possible that this hyperexcitability in response to injury could also be a risk factor for the development of future neurodegenerative disease. Indeed, a link has already been established between early adult hyperconnectivity in the gamma range (40-160 Hz) and an increased genetic risk of developing dementia (Koelewijn et al. 2019).

In the healthy brain, local cortical gamma oscillations are generated by mutual inhibitory loops within PV cell populations as well as inhibitory-excitatory loops between PV and pyramidal neurons (Buzsáki and Wang 2012). Given the critical involvement of PV interneurons in the generation of gamma-band activity (Cardin et al. 2009), the increase in specifically gamma band functional connectivity could suggest a circuit-level mechanism as the substrate of our observed hyperconnectivity. In addition to targeting distant-region pyramidal neurons, cortical pyramidal cells in layer 5 provide excitatory inputs to inhibitory VIP neurons (Modolo et al. 2020). These VIP neurons inhibit a target region’s SST neurons, and by doing so effectively disinhibit both pyramidal and PV neurons in that target region’s gamma generating circuit (Cottam, Smith, and Hausser 2013; Urban-Ciecko and Barth 2016; Xu et al. 2013; Muñoz et al. 2017; Karnani et al. 2016). In our model of bTBI, it is proposed that SST neuronal loss interferes with this mechanism and leads to a disinhibited flow of gamma oscillations to the target region, manifesting as increased functional connectivity in the gamma range. In addition, enhanced plasticity due to injury-related hyperexcitation could exacerbate this phenomenon by creating stronger synapses and thus more reliable transmission of the oscillations by pyramidal neurons (Cantu et al. 2015; Witkowski et al. 2019). Importantly, studies investigating other pathologies associated with an imbalance in excitation and inhibition such as Fragile X syndrome (Wang, Ethridge, et al. 2017), autism spectrum disorder (Wang et al. 2022) and schizophrenia (Di Lorenzo et al. 2015; Andreou et al. 2015) have reported the same finding of increased gamma connectivity. Thus, our results provide a potential mechanism mediating posttraumatic connectivity increases, which could in turn underlie cognitive symptoms and provide a therapeutic target for intervention.

When considering the impact of blast on electrophysiological measures such as functional connectivity, it is important to consider accompanying neuronal and structural changes. Changes in NeuN, a measure of overall neuron density, can be observed in more severe models of TBI such as experimental CCI models (Akamatsu and Hanafy 2020). However, in the current study, and consistent with previous research of single-event mild TBI (Ogino et al. 2022), NeuN staining indicated no significant change in overall neuron density in the regions of interest. Before drawing a conclusion of no neuronal loss, it must be considered that NeuN measurements have possible limitations when identifying interneuronal sub-type density changes. For example, a 40% loss of an interneuron population such as PV or SST, would only result in a ∼2% change in overall neuronal density when using NeuN. As a result, a lack of change in NeuN staining possibly only reflects the gross impact of blast on all neurons in a region. To counter this potential limitation of NeuN staining, we separately evaluated the neuronal density of two of the most abundant interneuron subtypes in the brain, PV and SST, which have been repeatedly implicated in TBI pathology (Cantu et al. 2015; Vascak et al. 2018; Masri et al. 2022; Frankowski et al. 2022). It has been further demonstrated that PV interneuron populations appear to be more vulnerable to nonblast induced TBI than SST interneurons (Nichols et al. 2018). In contrast to more severe models of TBI, we found SST interneurons to be more affected by a mild blast injury than PV interneurons. However, as with previous research, we found the 5^th^ layer to be most vulnerable to injury (Cantu et al. 2015).

When evaluating immunopositive staining results, conclusions regarding decreases in SST across all regions of interest contain an element of ambiguity, as they could be due to interneuron loss and/or downregulation of neuropeptide SST (which is not staining positive). For example, it has been shown in a more severe injury model that SST interneuron density decreases after non-blast TBI (Cantu et al. 2015). As with SST density, a decrease in immunopositively stained PV interneurons can indicate either PV interneuron loss or a reduction in the density of PV by these neurons (Filice et al. 2016). In this single blast model of mild TBI, only PV interneurons in the primary auditory cortex were affected, but this might be a result of hearing impairment or loss rather than the blast itself (Masri et al. 2022). This could be the case in our model since no ear protection was used in our experimental setup and could have led to temporary hearing loss due to eardrum rupture. Given that there was no interneuronal recovery over any time points post-injury, this could also suggest our results are showing interneuron cell death rather than down-regulation of their expression. Nevertheless, whether our results are an indication of interneuron loss or downregulation of the expression, or both, it could disrupt the E/I imbalance and cause neural hyperexcitability (Lodge, Behrens, and Grace 2009; Ferguson and Gao 2018).

In addition, our data show an immediate microglial density increase post-injury as a sign of inflammatory activation, which lasts up until 1-month post injury in the Au1 and RSC. Our results also demonstrate an increase in percentage area stained in all cortical regions other than the RSC at 3-months post-injury, indicating hypertrophy and potentially a change in phenotype. This result aligns with other acute time point studies of blast (Huber et al. 2016), and mild blast TBI studies that see both inflammatory responses at later time points (De Lanerolle et al. 2011; Goldstein et al. 2012; Masri et al. 2022; Ou et al. 2022). Finally, our results show that there was no significant increase in astrocytic activation at any time point after injury, thus suggesting no obvious astrocyte influence on the electrophysiological changes seen in our study. This is in contrast with previous work, where astrocytic activation is often reported, especially at acute timepoints post-injury (Sajja et al. 2012; Svetlov et al. 2010; Cernak et al. 2011). However, a study that investigated bTBI with a similar impulse energy and subject orientation to the blast wave obtained similar findings to ours (Rubio et al. 2021), possibly highlighting the importance of blast parameters in terms of determining the nature of injury outcomes.

In this study, we have developed a preclinical animal model of mild blast-induced TBI and demonstrated that the employed techniques and metrics of injury assessment reflect the subtle changes associated with mild injury. As electrophysiology is already used in the clinical setting, our employed outcome measures are highly translatable. We have combined histological findings only accessible in an animal model with clinically relevant EEG to obtain further clarity on the relationship between the structural and electrophysiological changes observed in TBI. We show the value of high-density EEG to assess injury, and therefore present this as a potential tool to be used for TBI diagnosis and management of treatment. Future research would do well to investigate the long-term structural causes of bTBI-induced hyperconnectivity and hyperexcitability to determine their contributions to neurogenerative disease.

## Acknowledgements

The authors would like to thank John Goodwin and Central Biomedical Services of Imperial College for their assistance, and Thuy-Tien Nguyen for help with the shock tube. We acknowledge the contribution made by Drs Dickinson and Campos-Pires in previous collaborative work using a blast model. We would also like to thank Dr Hannah Jones for creating the artificial rat used for shock tube optimisation.

## Funding

This study was funded through the Royal British Legion at the Centre for Blast Injury Studies, Imperial College London.

## Competing interests

The authors report no competing interests.

## Supplementary Information

### TRBL-CBIS shock tube

All blast experiments used the Royal British Legion Centre for Blast Injuries (TRBL-CBIS) shock tube at Imperial College London. The shock tube comprises three separate 1.22 m long tubes with an internal diameter of 59 mm that are bolted together with flanges. The first 1.22 m long tube acts as the driver section and connects to the following two tubes known as the driven section (Fig. 1A). The driver section was reduced to 7% capacity with use of a partitioning blanking plate and was separated from the driven section by replaceable 250 µm Mylar® diaphragms. This reduction was necessary to produce a Friedlander waveform (Supplementary Fig. 1A) (Nguyen, Wilgeroth, and Proud 2014; Nguyen et al. 2019; Nguyen et al. 2018). A shock wave was generated by filling the 7% driver section with compressed air until a Mylar® diaphragm burst, releasing the air through the driven section. The shock wave propagated down the driven sections and towards the right side of the rat head. Reproducibility of the shock wave was monitored using piezoelectric pressure sensors (Dytran Instruments 2300V1, California, USA) that were flush with the inside wall of the shock tube to give in-tube incident shock wave pressure readings from sensor 2 (Supplementary Fig. 1B). Control experiments were conducted using a phantom rat head model (as seen in Supplementary Fig. 1C), and a third sensor (sensor 3) was substituted for the rat’s head to characterise blast at this location.

### Blast Procedure

Due to logistical constraints related to the experimental procedures and COVID-19 restrictions, we were unable to randomise the rats for our experiments. The study involved a combination of blast traumatic brain injury (TBI) and EEG measurements, which required careful coordination between the experimental days and the availability of the animals and the experimenters. Given these time constraints and logistical challenges, randomising the animals at different time points, such as 1- month and 3-months, would have been highly inefficient and impractical for the study. To avoid confounding effects of skill level differences between experimentalists, each part of the procedure was performed by the same experimentalist across all experiments.

Animals were weighed and then anaesthetised in an induction chamber with 5% isoflurane at 2.5 L/min. Once anesthetised, an animal was moved to a physiological monitoring platform (Harvard Apparatus) where anaesthesia was maintained at 1.5-2.5% using a 2 L/min flow rate delivered via a polyethylene nose cone. Analgesia with buprenorphine (0.05 mg/kg s.c.; Vetergesic, UK) was administered 30 mins prior to blast, and lubrication was placed over the eyes. A closed loop temperature control system with a rectal probe was used to maintain body temperature at 37 °C. In addition, breathing rate and blood oxygenation were also monitored using the platform. Once an animal was stable under anaesthesia (approx. 20 mins), it was moved into a protective shield on a platform fastened to the shock tube exit. The animal was placed perpendicular to the shock tube exit, with the snout positioned 4.5 cm from the shock tube exit (with right side of the animal closest to blast) and 5cm from the exit of the protective shield (schematic in Fig. 1A). This placement allowed the body to be off-centre to the tube exit to prevent blast lung injury. Blood oxygenation continued to be monitored whilst the animal was on the blast platform and breathing rate was manually counted prior to blast. The animal was fixed from the torso down in the prone position inside the protective tube whilst the head was free to move in response to the shock wave. Anaesthesia was delivered via the soft polyethylene nose cone which would then be expelled during blast. After blast exposure the animal was immediately removed from its protective shield and placed on a heat mat in a lateral recumbent position. Oxygen-enriched air was provided, and blood oxygenation levels were monitored during recovery. 3 mL of sodium chloride 0.18% and glucose 4% saline solution for infusion (Baxter) was administered. Blast characteristics are presented in supplementary Table 1. Loss of righting reflex (LORR) was measured from the time of the blast until an animal had all four paws touching the ground in the recovery cage. The loss of righting reflex was used as a proxy for loss of consciousness (LOC) as used in humans.

Once an animal had fully recovered, it was returned to an individual cage and monitored until the end of the experimental day when it was returned to a cage of four. Animals were given buprenorphine (mixed in Hartley’s strawberry, 1:2 jelly/water, 0.3 mg/kg p.o.) and weighed twice a day for 3 days. This was followed by daily welfare checks based on a based on a scoring system (Morton and Griffiths 1985) and weighing once a day for two weeks, and then once-a-week welfare checks until termination. Typical signs of TBI should develop on recovery from anaesthesia and over the subsequent 72 hours, a period covered by the monitoring scheme described above. Animals were considered to have mild TBI due to ∼30 min LORR (Supplementary Table 1) and a Morton and Griffiths welfare score of below 2, indicating no signs of pain, distress or weight-loss post-injury, together with no pulmonary blast lung damage induced mortality.

Humane endpoints were set as follows: Any animal experiencing a high level of distress or a reduction in body weight of or more than 20% would be humanely killed by a Schedule 1 method. If an animal displayed 2 or more of the following clinical signs, it would be killed by a Schedule 1 method: piloerection, hunched posture, reduced mobility, pall or, ocular or nasal discharge.

For the acute electrophysiological cohort, blast-exposure was identical to the chronic cohorts except for the anaesthetic protocol. More specifically, animals were anaesthetized using urethane as described in the ‘Electrophysiological Recordings – Surgery’ section of the supplementary methods. This anaesthetic protocol results in a stable anaesthetic depth and thus after baseline recordings of brain activity, animals were placed at the shock tube exit for the injury procedure. After injury, animals were transferred to a different room for post-injury recordings while still under terminal general anaesthesia.

### High speed videos

High-speed videos were used as quality control to ensure the animals were receiving the blast in the same manner similar to Mishra et al. (Mishra et al. 2016). (Software used: Physlets Tracker 5.1.5 (project source of: Open Source Physics www.compadre.prg/osp)).

### Tissue processing

PFA-fixed brains of the animals were blocked with a 3D printed brain matrix – and split into 9 2 mm blocks of tissue. All tissue processing was performed by the same experimenter. The block containing the regions of interest were embedded in paraffin and serially sectioned (7 mm) on a microtome. Slices were mounted onto individual slides, with 3 slices used per animal spanning ∼100 µm.

### Immunostaining

All immunostaining and analysis procedures were conducted by same investigator.

### GFAP and NeuN

Sections were deparaffinised and rehydrated using xylene and decreasing concentrations of ethanol. The tissue was washed in 1x PBS permeabilised and endogenous peroxidase quenched using 1% Hydrogen Peroxidase (H2O2) in 1x PBS + 1% Triton-100-X (PBS-Tx) (w/v) for 30 mins, followed by another 1x PBS wash and then antigen retrieval in 0.01M Citrate Buffer (pH6) for 20 mins in a steamer. Sections were incubated overnight at 4°C with the primary antibody GFAP (polyclonal, Rabbit IgG, #Z0334, DAKO, Denmark, 2.9 g/L, 1:2000) or NeuN (polyclonal, mouse IgG, #MAB377, Millipore, 1 mg/mL, 1:1000).

On the following day Super Sensitive polymer-HRP (BioGenex) IHC detection system was applied according to manufacturer’s protocol. The slides were then stained with 3,3′Diaminobenzidine (DAB,GFAP: 2 min, NeuN: 90 seconds) dehydrated, and cover slipped.

### PV and IBA1

Tissue preparation (deparaffinisation, rehydration, endogenous peroxidase quenching, antigen retrieval) was the same as described before. This was followed by blocking using 10% foetal bovine serum (FBS), 10% Normal Goat Serum (NGS) and 1% Bovine Serum Albumin (BSA) for 2 hours. Slides were incubated overnight at room temperature with the primary antibody IBA1 (polyclonal, rabbit IgG, Fujifilm Wako Pure Chemical Corporation, #019-19741, 1:1000) and overnight at 4°C PV (polyclonal, rat IgG, #PV-235, Swant, 1:10,000) in 2%FBS/2%NGS/0.2%BSA PBS + 1% Triton-100-X (PBS-Tx) (w/v).

The following day, slides were incubated with the biotinylated secondary antibody (goat antirabbit biotinylated, #BA-1000, Vector Labs, 1:250, or Goat anti-rat biotinylated #BA-9400, Vector Labs, 1:250) in 2%FBS/2%NGS/0.2%BSA PBS + 1% Triton-100-X (PBS-Tx) (w/v) (table 4) for 90 mins. The slides were then incubated with avidin-biotin complex (ABC) solution in PBS for 40 mins and then stained with DAB (IBA1: 4 mins and PV: 2 mins). For IBA1, the slides were dehydrated and coverslipped as previously described. For the PV they were counterstained with haematoxylin (#HEMM-35/21, Solmedia) for 20 seconds before dehydration and coverslipping.

### SST

Tissue preparation was the same as described before, followed again by tissue blocking using 10% foetal bovine serum (FBS), 10% Normal Horse Serum (NHS), and 1% Bovine Serum Albumin (BSA) in 1% Triton-100-X (PBS-Tx) (w/v) for 2 hours. Slides were then incubated overnight at 4°Cwith the primary antibody SST (purified recombinant rabbit monoclonal, rabbit IgG, #ZRB1042, EMD Millipore Corporation, 1:100) in 2%FBS/2%NGS or 2% NHS/0.2%BSA + 1% Triton-100-X (PBS-Tx) (w/v).

On the following day slides were incubated with the ImmPRESS kit (#MP-7401, ImmPRESS horse anti-rabbit IgG polymer kit, peroxidase, 1-2 drops/brain) for 30 mins. The slides were then stained with DAB for 2 mins.

### NFL Immunofluorescence (IF) staining

Sections were deparaffinised and rehydrated, then subjected to antigen retrieval. Following this, slides were blocked with 10% FBS/10%NGS/1%BSA + 1% Triton-100-X (PBS-Tx) (w/v) for 2 hours. The slides were then incubated with the primary antibody NFL (polyclonal, rabbit IgG, #171-002, 1:10,000) overnight at 4 degrees. After overnight incubation the sections were treated with the secondary antibody (Goat anti-rabbit, #A21245, Invitrogen Alexa Flura 647, 1:200) in 2%FBS/2%NGS or 2% NHS/0.2%BSA 1% + 1% Triton-100-X (PBS-Tx) (w/v) (table 4) and protected from light for 90 mins. Following incubation, the slides were incubated with 4′,6diamidino-2-phenylindole (DAPI) 1:1000 for 5 mins then and mounted with Prolong Diamond Antifade (Life technologies, #P36970) then sealed with all-purpose nail varnish and cured overnight at 4 degrees.

### Immunostaining image Analysis

Regions of interest (ROIs) were selected based on different cortical column orientations relative to the incident shockwave. The ROIs were primary auditory cortex (Au1), primary visual cortex (V1), retrosplenial cortex (RSC), and the CC (Fig. 1D). Density of NeuN positive neurons was assessed individually in each cortical layer. In all ROIs, a box was drawn to span the cortex from the bottom of layer 1 to the bottom of layer 6. Box width was set at 700 µm for the Au1, 400 µm for the RSC, and 500 µm for the V1 cortex. Boxes were divided into cortical layers and analysis was performed for both hemispheres. Each layer per region of interest was analysed separately. Neuronal density was calculated using a HALO® cytonuclear algorithm and presented as neuronal density cell/mm^2^. For the NeuN analysis in the Au1 3 animals at 1-month post-injury were excluded due to tearing or staining artefacts in that region. For the V1 and RSC, 1 animal was not included at 7-days post-injury, and 3 animals at 1-month post-injury were excluded for the same reasons. The same regions of interest were used for GABAergic inhibitory interneurons analysis, but cells were counted manually.

Results were presented as cellular density (cell/mm^2^). For the Parvalbumin staining in the Au1, and the RSC, 1 animal was not included for the analysis due to staining artefacts in the tissue at 1month post-injury. At 1-month post-injury 2 animals were excluded for the same reason in the V1. For astrocytic analysis, regions of interest had an overall box size comparable to those of the neuronal stains, e.g., 700 µm box width for Au1 cortex spanning from pia to white matter. The CC was outlined for analysis to be compared against the Naïve/Sham group. Using a HALO® algorithm for % area covered, a threshold between background and positive staining was established and used to calculate % area stained. For the RSC and V1, 2 animals were excluded from the 7-days post-injury group and 3 animals from the 1-month post-injury group due to tearing or staining artefacts. For the CC 1 animal was excluded from the 1-month post-injury group for similar reasons. Microglial density quantification was performed using a HALO microglial algorithm. For all regions 2 animals were excluded from the 7-days and 1-month post-injury group due to staining artefacts. Microglial % area stained was calculated in a similar to that of the astrocytes. IBA1 % area stained had 4 animals excluded from the Naïve/Sham group due staining artefacts and 2 excluded from the 7-days post-injury group, for the same reasons. Neurofilament light (NFL) was analysed in the corpus callosum using HALO’s fluorescent intensity covered algorithm and intensity was normalised to two small low fluorescent areas of the periaqueductal grey (PAG). 1 animal was excluded from the 1-month and 3-months post-injury groups due to NFL staining artefacts. The data were presented as NFL intensity (% change from Naïve/Sham). Luxol fast blue (LFB) CC was used to measure CC thickness and averaged per animal, then normalised to the Naïve/Sham group and presented as % of Naïve/Sham. 1 animal was excluded from the 6 Hr, 1-month and 3-months post-injury groups due to tearing.

### Electrophysiological Recordings

#### Surgery

Surgery was performed by the same experimenter to ensure consistency. Induction was performed using isoflurane (5% concentration, 2 L/min flow rate) inside an induction chamber, followed by stabilisation of respiratory rate using a face mask and titrating the concentration to a target of 80 breaths/minute. This was followed by an intraperitoneal injection of urethane (2.7 ml/kg, 1.35 g/ml, Sigma-Aldrich). Follow-up doses of 10% of the initial dose, were administered as necessary to ensure loss of eye-blink and paw-withdrawal reflexes. Atropine (0.66 ml/kg, 1% w/v) was administered subcutaneously to reduce mucous secretions.

After a minimum 1hour period, the scalp of the animal was shaved, and bupivacaine (7.5 mg/kg, Marcaine) was administered subcutaneously to the area for local analgesia. An incision was made to expose the skull, and tissue was cleared from the skull using blunt dissection. Throughout the anaesthetic induction and surgery, blood oxygen saturation and respiratory rate was monitored and normothermia was achieved using a physiological monitoring platform with closed-loop temperature control (Harvard Apparatus). Then throughout the course of the experiment, temperature was controlled using isothermal pads (Fisher Scientific) to prevent the introduction of electrical noise to the electrophysiology. Anaesthetic depth was frequently monitored by means of testing eye-blink and paw-withdrawal reflexes.

#### Recording Setup

The multielectrode EEG array (Neuronexus) used 500 μm diameter Platinum electrodes on a 15 μm thick Polyimide substrate, spanning 11.7 mm on the rostro-caudal and 9.6 mm on the left- toright axis, with a minimum inter-electrode distance of 1.2 mm. Electrode impedance was below 10 kOhm at 1 kHz.

Signals were digitized proximally to the animals’ heads using a Neuronexus SmartLink^TM^ headstages. The reference signals were obtained from copper wires fitted through and wound around surgical needles inserted into the nuchal musculature. All signals were analog filtered between 1.1 Hz and 15 kHz and subsequently sampled at 30 kHz using a Neuronexus Smartbox Pro^TM^ acquisition board (16-bit A/D converter) interfacing with the Radiens Allego^TM^ acquisition software (Neuronexus) running on a Windows computer and saved to disk.

#### Electrophysiological Data Preprocessing

Data were preprocessed by the same investigator to ensure consistency. Rejection of line noise with the CleanLine method (Bigdely-Shamlo et al. 2015) was performed in two steps with different parameter values for the sliding window – 4 s windows with 50% overlap and 25 s windows with no overlap – that we empirically found resulted in satisfactory suppression of line noise. All other parameters were set at their default values. For the Artifact Subspace Reconstruction algorithm, we used a value of 12 for the burst criterion and 0.6 for the correlation criterion, with all other parameters set at default values. Datasets were excluded from further analysis in cases where line noise was not effectively suppressed, or the automated artifact rejection procedure led to more than 20% of the datapoints being rejected. 1 animal in the blast 1-month group and 2 animals in the sham 3-months group were excluded based on these criteria.

### Statistical Analysis

For the electrophysiological results, statistical analysis was performed using Matlab^TM^. For the acute electrophysiological cohort’s global power and connectivity comparisons, we performed a permutation based paired *t-*test using functions from the Matlab^TM^ Statistics and Machine Learning Toolbox (function *ttest*). For the chronic electrophysiological cohort’s power comparisons at both the global and electrode/electrode pair level and for comparisons of global connectivity, a permutation based 2-way ANOVA (factor 1: injury group (blast vs sham), factor 2: timepoint (1month post-blast vs 3-months post-blast, plus interaction term) test was performed (Anderson 2001), where *P*-values were calculated by comparing the observed F-values to the 95^th^ percentile of a distribution of statistics obtained under a permutation of group labels (10,000 permutations). In the case of a significant interaction, post-hoc permutation-based independent samples Welch’s unequal variances t-test were performed by adapting functions from the Matlab^TM^ Statistics and Machine Learning Toolbox (function *ttest2*) between the pairs of interest (Blast vs Sham at 1month, Blast vs Sham at 3-months and Blast at 1-month post-injury vs Blast at 3-months postinjury). To correct for multiple comparisons, the Bonferroni method was used for measures of global power and connectivity (*n* = 6 frequency bands), electrode-specific power for each band (*n* = 32 electrodes) and pairwise *t*-tests (*n* = 3 pairs). To assess differences at the level of individual edges, the timepoint factor was collapsed and an independent samples *t*-test was performed using the network-based-statistic toolbox (Zalesky, Fornito, and Bullmore 2010). Correction for multiple comparisons was performed using the toolbox implementation of the False Discovery Rate method with the corrected *P*-threshold set at *P* = 0.05 and 50000 permutations.

The histological data was analysed in GraphPad Prism (9.4.1). The control group consisted of naïve and sham pooled data as there were no significant differences between them. NeuN, PV, and SST cell densities were analysed per cortical layer for each region of interest. For these analyses, a Shapiro-Wilk normality test was performed followed by a one-way ANOVA (factor: time (Naïve/Sham, 6-hours, 7-days, 1-month post-blast vs 3-months post-blast)). Multiple-comparison tests were conducted for each region’s layers using a Two-stage linear step-up procedure of Benjamini, Krieger and Yekutieli (BKY) False Discovery Rate (FDR) approach with *Q* = 5%. Post-hoc comparisons (Uncorrected Fisher’s Least Significant Difference (LSD)) were performed against the Naïve/Sham cohort. To investigate glial changes, the analysis was performed for each region of interest. A Shapiro-Wilk normality test was performed followed by a one-way ANOVA. To investigate glial activation due to blast in the CC a Shapiro-Wilk normality test and a KruskalWallis test was performed. Neurofilament light data underwent normality tests followed by a oneway ANOVA. For each histological analysis, the data underwent an outlier identification check using the ROUT method provided by Prism (Q=1%). If a value was identified as an outlier, it was excluded from the dataset. Data were presented as ± SEM unless stated otherwise, and the threshold for significance was set at *P* = 0.05.

Investigators were not blinded to the outcomes assessments. No sample size or effect size calculations were performed.

## Supplementary Figures and Tables

**Supplementary figure 1.**
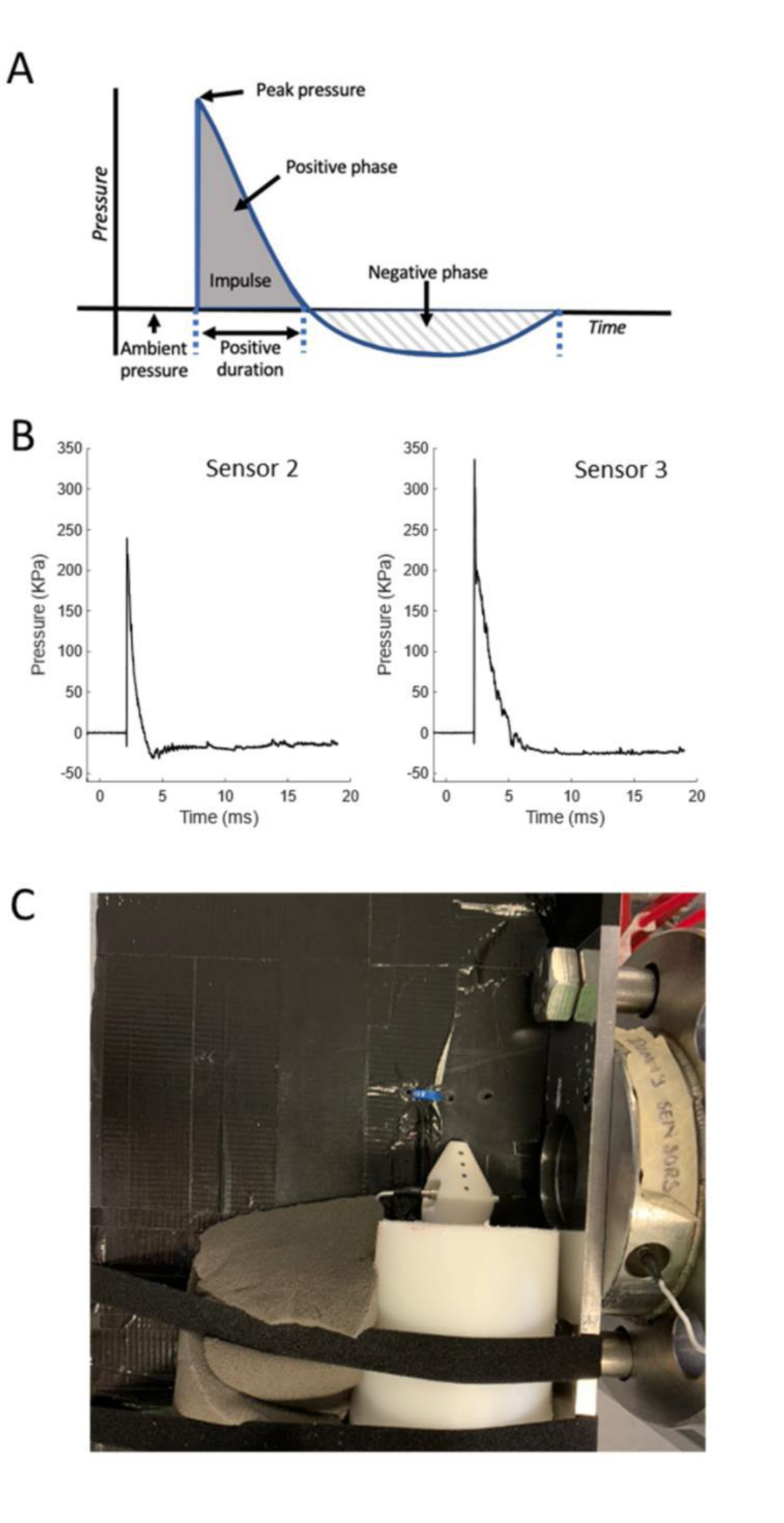
**(A)** Idealised Friedlander waveform. **(B)** Representative pressure-time graphs of the shock wave from, left graph; incident pressure sensor 2 and right graph; total pressure sensor 3. **(C)** 3^rd^ Sensor 3D rat. 3D printed rat secured in place with hole screwed into the head for the sensor to sit flush with the edge of the “rat head” for accurate measurement of total pressure from the oncoming blast wave.

**Supplementary figure 2.**
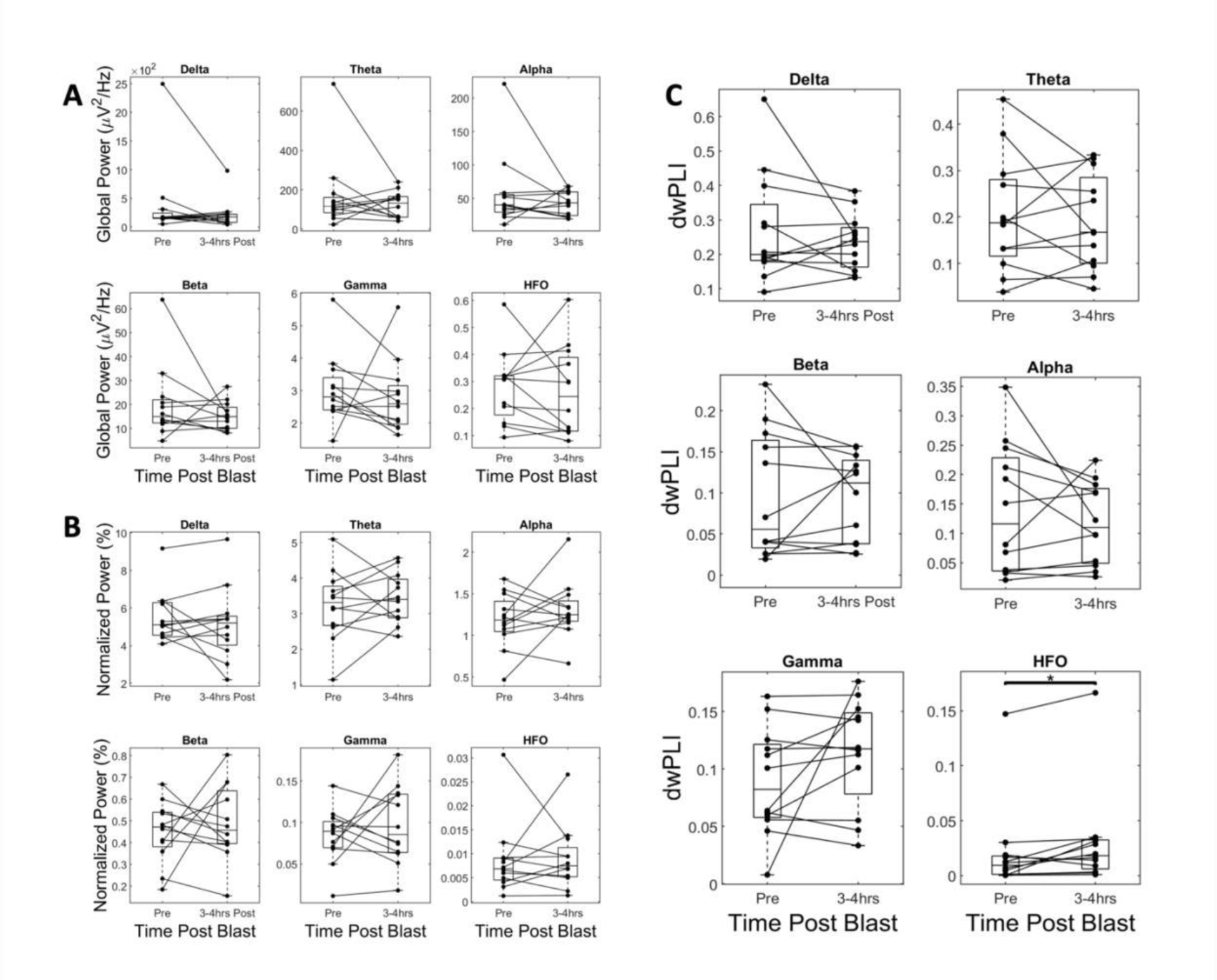
IBA1+ cell density in all regions of interest over reported timepoints (A-E) IBA1+ staining of the auditory cortex. **(F-J)** IBA1+ staining in the retrosplenial cortex. **(KO)** IBA1+ staining in the primary visual cortex (V1). **(P-T)** IBA1+ staining in the corpus callosum. **(A/F/K/P)** Naïve/Sham, **(B/G/L/M)** 6 Hr post-injury, **(C/H/M/R)** 7-days post-injury, **(D/I/N/S)** 1month post-injury, **(E/J/O/T)** 3-months post-injury. **(U-X)** quantification of IBA1+ for each region of interest using one-way ANOVA (Au1: Naïve/Sham: n = 13, 6 Hr: n = 6, 7-day: n = 4, 1-month: n = 6, 3-months: n = 6. V1: Naïve/Sham: n = 13, 6 Hr: n = 6, 7-day: n = 4, 1-month: n = 6, 3months: n = 6. RSC: Naïve/Sham: n = 13, 6 Hr: n = 6, 7-day: n = 4, 1-month: n = 6, 3-months: n = 6. CC: Naïve/Sham: n = 13, 6 Hr: n = 6, 7-day: n = 4, 1-month: n = 6, 3-months: n = 6.

**Supplementary figure 3.**
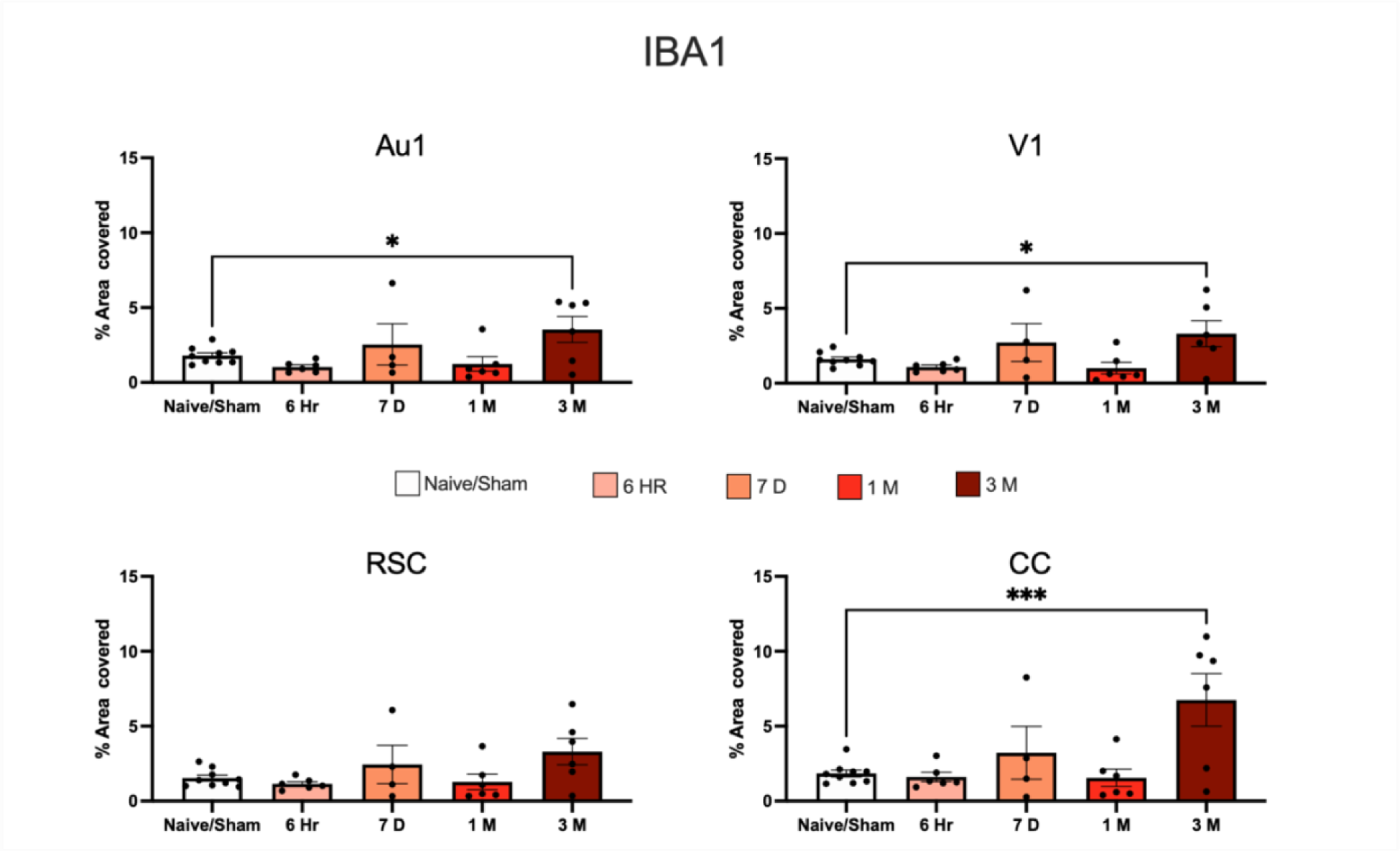
Analysis of IBA1 % area covered in all regions of interest at (6 Hr, 7- days post-injury, 1-month and 3-months post-injury) in the primary auditory cortex (Au1), retrosplenial cortex (RSC), primary visual cortex (V1) and the corpus callosum (CC). One-way ANOVA revealed significant increase in % area stained at 3-months. Quantification of IBA1+ for each region of interest using one-way ANOVA data presented mean ± SEM, \**P*<0.05 (Au1: Naïve/Sham: n = 9, 6 Hr: n = 6, 7-day: n = 4, 1-month: n = 6, 3-months: n = 6. V1: Naïve/Sham: n = 9, 6 Hr: n = 6, 7-day: n = 4, 1-month: n = 6, 3-months: n = 6. RSC: Naïve/Sham: n = 9, 6 Hr: n = 6, 7-day: n = 4, 1-month: n = 6, 3-months: n = 6. CC: Naïve/Sham: n = 9, 6 Hr: n = 6, 7-day: n = 4, 1-month: n = 6, 3-months: n = 6).

**Supplementary figure 4.**
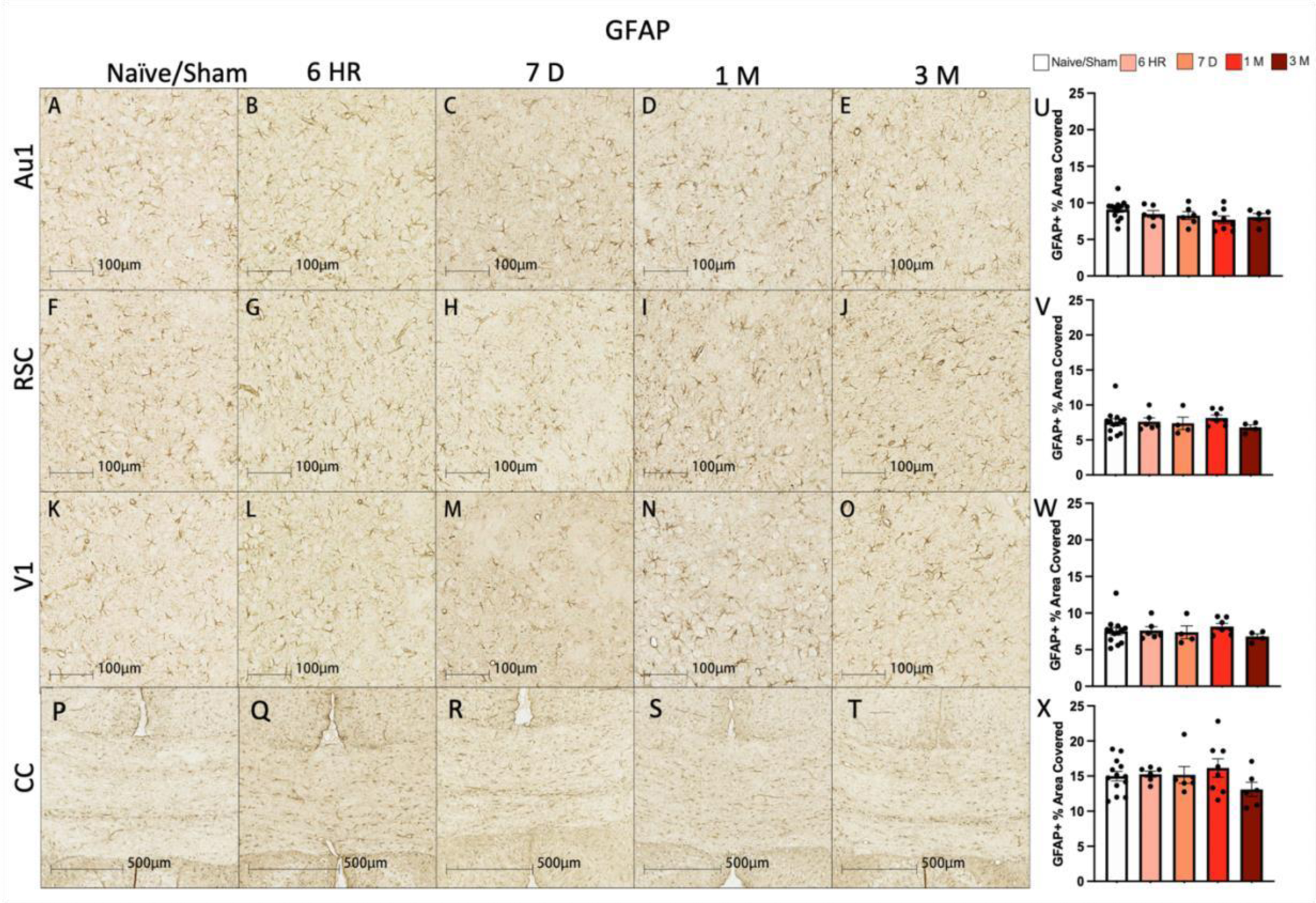
GFAP % area covered in all regions of interest. (A-E) GFAP+ % staining in of the auditory cortex. **(F-J)**, retrosplenial cortex **(I-O)** GFAP staining in the primary visual cortex (V1) cortex. **(P-T)** GFAP staining in the corpus callosum (CC). **(A/F/K/P)** Naïve/Sham, **(B/G/L/M)** 6 Hr post-injury, **(C/H/M/R)** 7-days post-injury, **(D/I/N/S)** 1-month post-injury, **(E/J/O/T)** 3-months post-injury. **(U-X)** quantification of IBA1+ for each region of interest using one-way ANOVA data presented mean ± SEM. \**P*<0.05, \*\**P*<0.01, \*\*\**P*<0.001 and \*\*\*\**P*<0.0001 (Au1: Naïve/Sham: n = 13, 6 Hr: n = 6, 7-day: n = 6, 1-month: n = 8, 3-months: n = 6. V1: Naïve/Sham: n = 13, 6 Hr: n = 6, 7-day: n = 4, 1-month: n = 5, 3-months: n = 6. RSC: Naïve/Sham: n = 13, 6 Hr: n = 6, 7-day: n = 4, 1-month: n = 5, 3-months: n = 6. CC: Naïve/Sham: n = 13, 6 Hr: n = 6, 7-day: n = 5, 1-month: n = 8, 3-months: n = 6)

**Supplementary figure 5.**
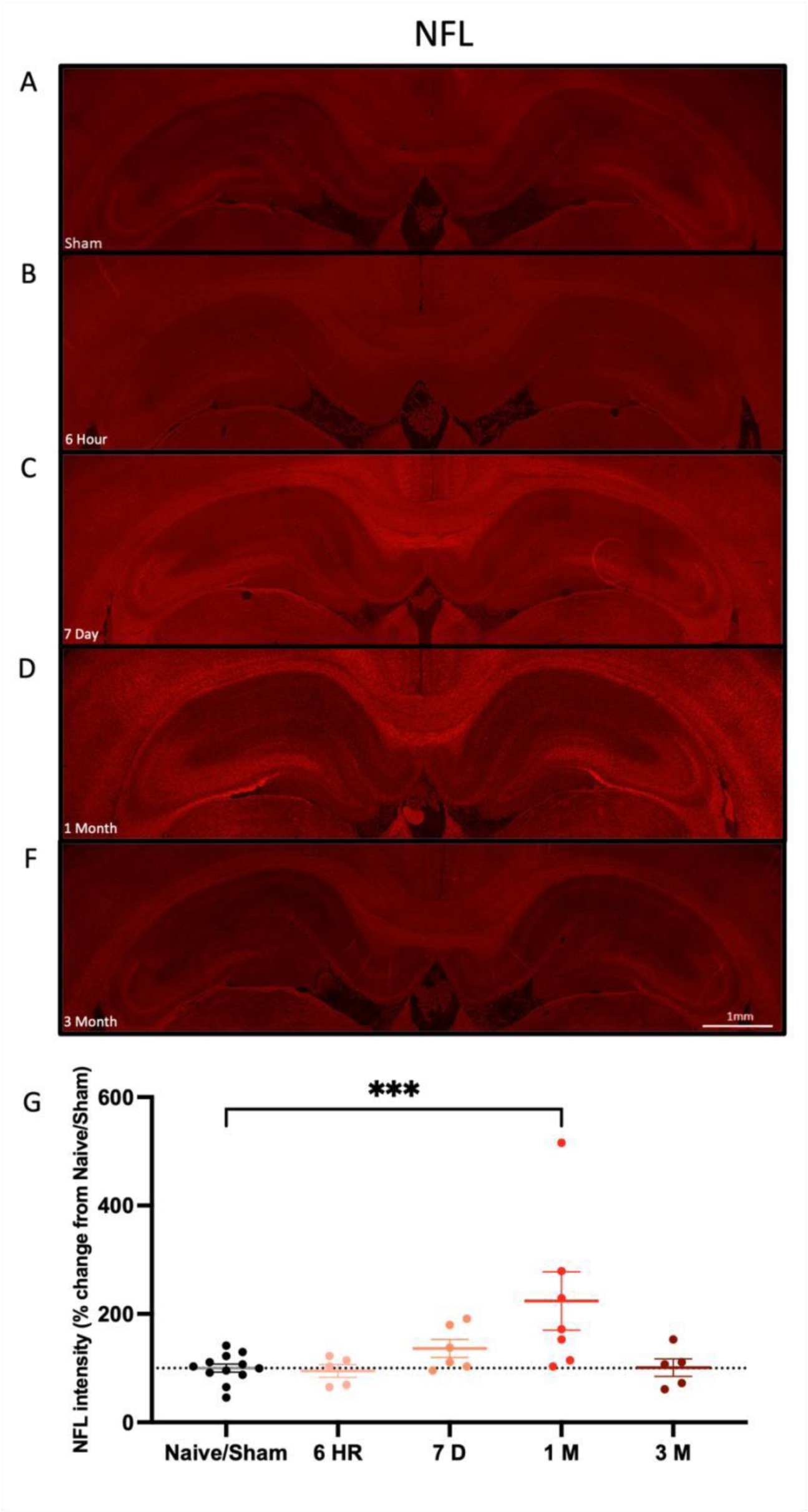
Neurofilament light fluorescence intensity in the corpus callosum (A-F) Representative staining of the corpus callosum of Alexa 546 of NFL of all time points postinjury. **(G)** Quantification of Alexa 546 immunofluorescence intensity for neurofilament staining in the whole of the corpus callosum; quantified using one-way ANOVA. Data presented mean ± SEM. \**P*<0.05, \*\**P*<0.01, \*\*\**P*<0.001 and \*\*\*\**P*<0.0001 (Naïve/Sham: n = 13, 6 Hr: n = 6, 7day: n = 6, 1-month: n = 7, 3-months: n = 5).

**Supplementary figure 6.**
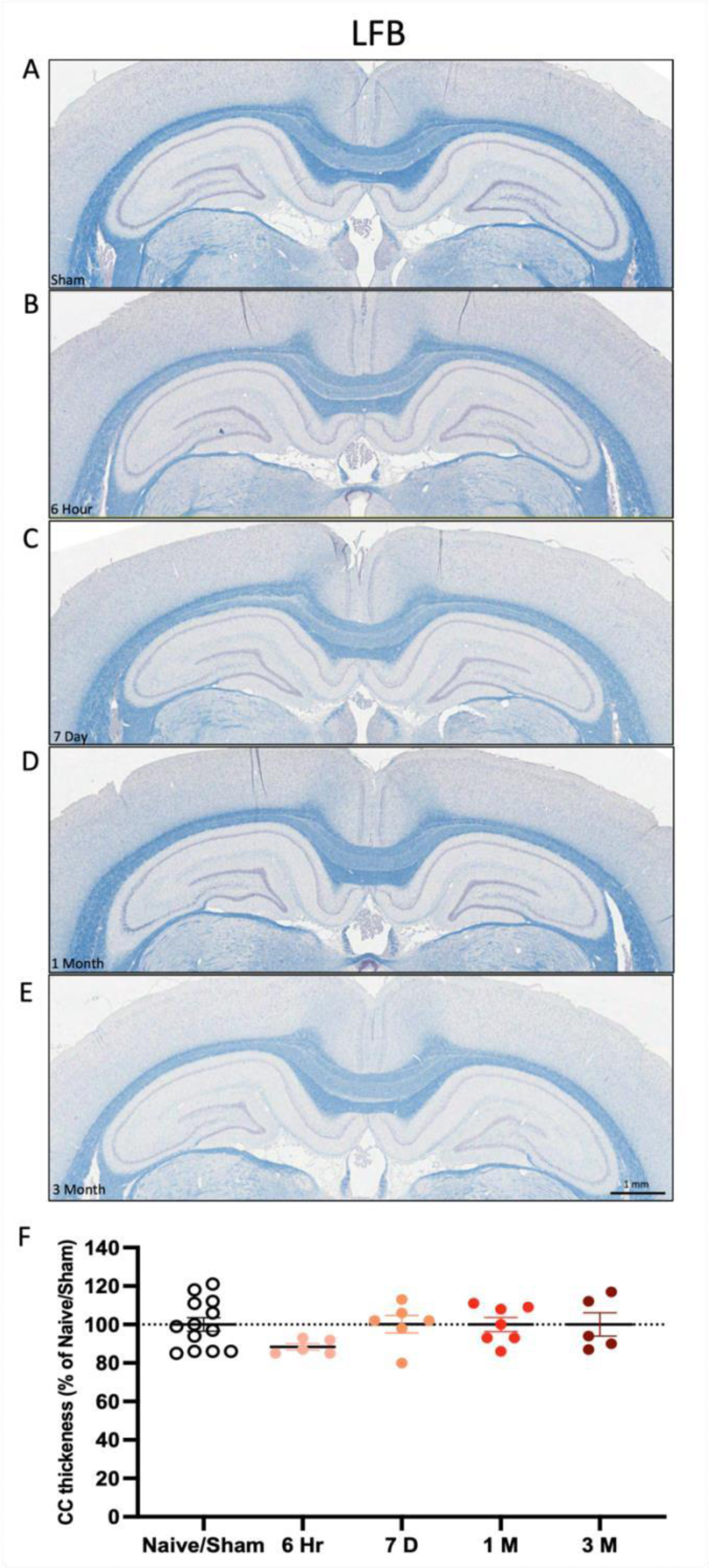
Luxol fast blue staining of white matter tract. **(A-E)** representative half-brain micrographs of Luxol fast blue staining from Naïve/Sham to 3-months post-injury. **(F)** quantification of 5 separate segments normalised to the contralateral hemisphere and then compared against Naïve/Sham data presented mean ± SEM. \**P*<0.05, \*\**P*<0.01, \*\*\**P*<0.001 and \*\*\*\**P*<0.0001 (Naïve/Sham: n = 13, 6 Hr: n = 5, 7-day: n = 6, 1-month: n = 7, 3-months: n = 5).

**Supplementary Table 1:**
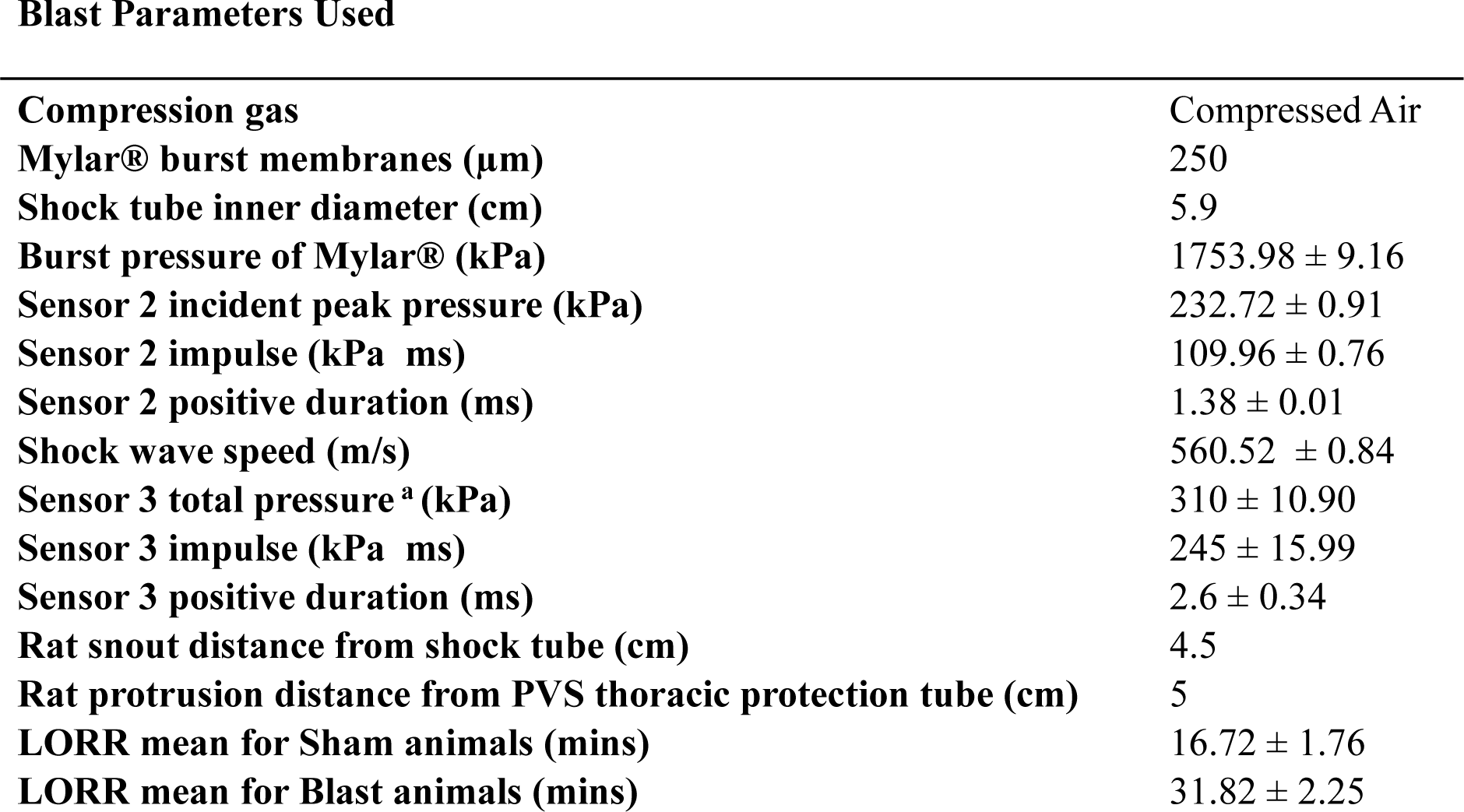
Blast characteristics. ^a^ Total pressure is the combination of both incident and reflected pressure is presented as mean ± SEM from 4 trials obtained during testing what the animal would receive. The total pressure is from 4 cm from the shock tube exit.

## Notes

### Competing Interest Statement

The authors have declared no competing interest.

